# Cross-species machine learning improves diagnostic classification of human psychiatric disorders

**DOI:** 10.1101/812693

**Authors:** Yafeng Zhan, Jianze Wei, Jian Liang, Xiu Xu, Ran He, Trevor W. Robbins, Zheng Wang

## Abstract

Psychiatric disorders often exhibit shared (co-morbid) symptoms, raising controversies over accurate diagnosis and the overlap of their neural underpinnings. Because the complexity of data generated by clinical studies poses a formidable challenge, we have pursued a reductionist framework using brain imaging data of a transgenic primate model of autism spectrum disorder (ASD). Here we report an interpretable cross-species machine learning approach which extracts transgene-related core regions in the monkey brain to construct the classifier for diagnostic classification in humans. The cross-species classifier based on core regions, mainly distributed in frontal and temporal cortex, identified from the transgenic primate model, achieved an accuracy of 82.14% in one clinical ASD cohort obtained from Autism Brain Imaging Data Exchange (ABIDE-I), significantly higher than the human-based classifier (61.31%, *p* < 0.001), which was validated in another independent ASD cohort obtained from ABIDE-II. Such monkey-based classifier generalized to achieve a better classification in obsessive-compulsive disorder (OCD) cohorts, and enabled parsing of differential connections to right ventrolateral prefrontal cortex being attributable to distinct traits in patients with ASD and OCD. These findings underscore the importance of investigating biologically homogeneous samples, particularly in the absence of real-world data adequate for deconstructing heterogeneity inherited in the clinical cohorts.

**One Sentence Summary:** Features learned from transgenic monkeys enable improved diagnosis of autism-related disorders and dissection of their underlying circuits.

## Introduction

Comparative analyses of brain connectomics in closely related species can shed light on changes of structural and functional network architecture occurring during evolution (1–5), which provides crucial insights for the development of viable diagnostic biomarkers for brain disorders. The pursuit of imaging-based biomarkers for autism spectrum disorder (ASD) has been challenged by a lack of biological accounts (6, 7) and a mix of divergent reports (8–10), mainly constrained by its substantial heterogeneity and co-morbidity with other psychiatric disorders such as obsessive-compulsive disorder (OCD) and attention deficit–hyperactivity disorder (ADHD) (11–14). Cumulative evidence has demonstrated the feasibility of constructing genetically engineered non-human primates to deepen mechanistic understanding of psychiatric disorders and development of more effective therapeutics (15–18). Despite its great potential, there exists no such roadmap for develop methodologies for improving cross-species translational research from direct cross-species extrapolation from monkey results to cross-species translational mapping in the field thus far (19).

To this end, we proposed a novel cross-species machine learning framework that leverages connectome-based features learned from a primate genetic model of ASD and then built a classifier for diagnostic utility in humans. We began by postulating what features can be learned from the monkey model that captures the underlying neural pathophysiology likely shared with human ASD patients. Within the network graph setting (4), two basic elements of brain circuitry: nodes (brain regions) and interconnecting edges (connections between pairs of nodes) are considered (1, 4, 20). Our intuition was that characteristics of nodes are more likely to be evolutionarily conserved between the primate species, given substantial variations of edges subserving species-specific behavioral and cognitive adaptations (1, 4). Moreover, supposing that only a subset of brain regions (not the entire brain) are particularly relevant to the core neuropathology of autism (termed as “core regions” thereafter) (9, 21), we deduced that only these “core regions” are useful for greatly reducing the complexity of cross-species mapping of this framework (Fig. 1A).

**Fig. 1.**
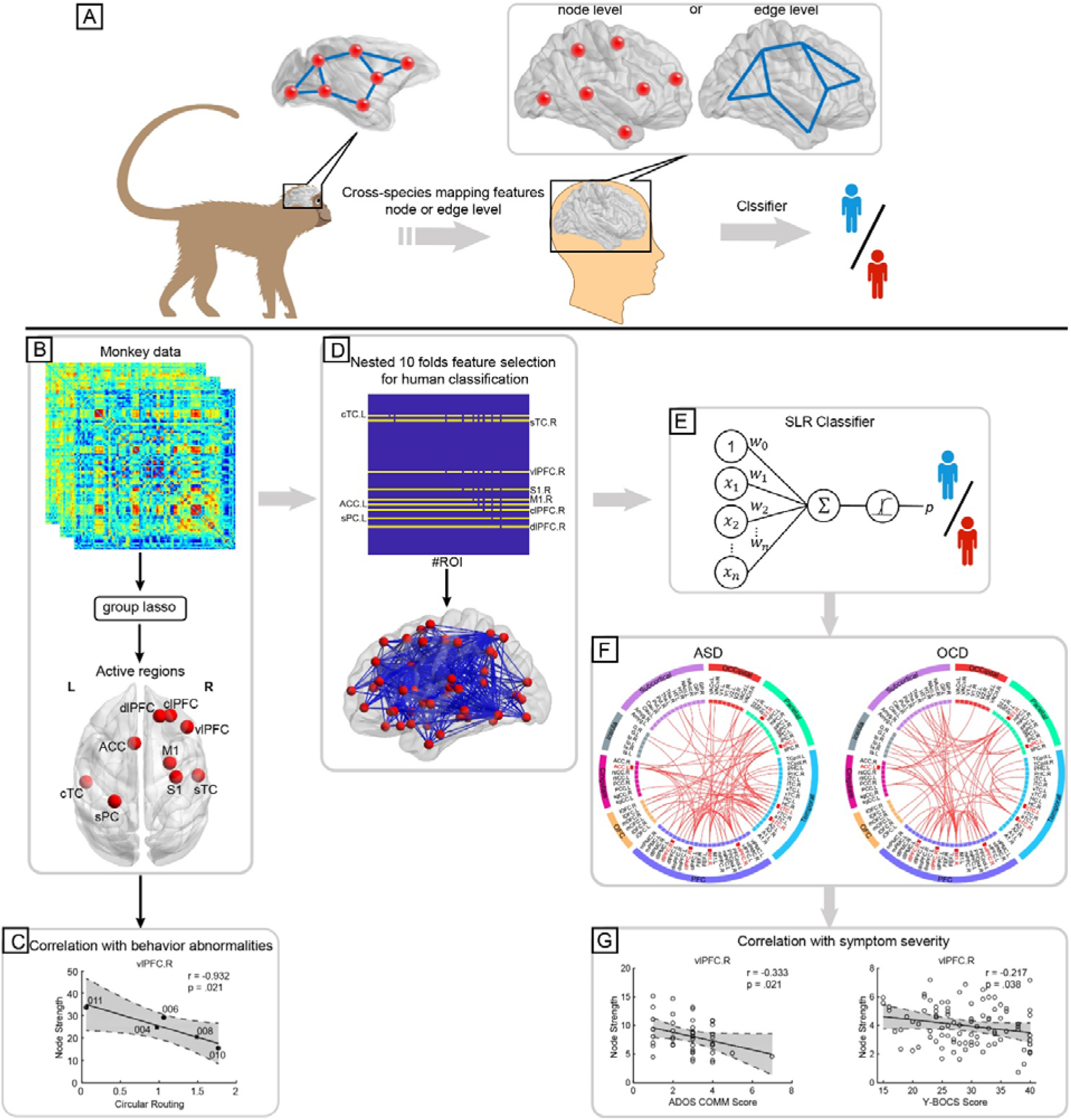
Schematic of a cross-species machine learning framework. (A) Illustration of a cross-species circuit mapping strategy, which is formulated on the basis of the intuition that characteristics of the brain circuitry between nonhuman and human primates are evolutionarily conserved at the node or edge level. (B) Using a group lasso to identify core regions in transgenic monkeys (C) Associations between characteristics of core regions and behavior abnormalities in transgenic monkeys. (D) Feature engineering in the human data is initiated by a one-to-one projection of core regions from monkey to human and then subjected to a nested 10-fold feature selection for use in human classification. (E) Optimal features were input into the sparse logistic regression (SLR) procedure for human classification. (F) Spatial profiles of the identified FCs in the ASD and OCD classifiers. (G) Associations between characteristics of nodes and edges in the ASD and OCD classifiers and symptom severity in patients.

Using whole-brain functional connectivity datasets obtained from wild-type and human Methyl-CpG binding protein 2 (MECP2) transgenic monkeys (17), and from multicenter ASD databases, available from the Autism Brain Imaging Data Exchange (ABIDE-I, N = 1112; ABIDE-II, N = 1114) (8, 22), OCD (N = 186) (23), and ADHD publicly available from ADHD-200 (N = 776) (24), we aim to test two specific hypotheses: (i) whether the monkey-based features (core regions) can improve the diagnosis accuracy of the ASD cohort; (ii) whether these monkey-based features can be generalised to improve the discriminative classification of OCD and ADHD from their healthy controls (HCs). To further parse the biological accounts of the monkey-based classifiers, stepwise linear regression models based on these classifiers are built to predict clinical measures of symptom severity of psychiatric patients (Fig. 1).

## Results

A total of 144 fMRI datasets from 5 *MECP2* transgenic (TG) and 11 wild-type (WT) monkeys were collected. We also analyzed ASD data from ABIDE-I/II repository (8, 22) and ADHD cohort from ADHD-200 (24), two publicly available multisite datasets of resting-state functional imaging data, and one OCD cohort from our institutional database (25).

### Core regions mapping between primate species

In order to determine the core regions in the monkey datasets for use in cross-species mapping, we adopted the sparse linear regression model based on the group lasso penalty (26, 27) which selects groups of variables (regions) based on predefined variable groups (all edges connected to one region being treated as a group, i.e. rows in an adjacency matrix). The penalty is deployed to regularize the problem to impose a network structure, i.e., core regions versus non-relevant regions here (Fig. 1B, see Materials and Methods for details of the group lasso method). A total of 144 fMRI datasets from 5 *MECP2* transgenic (TG) and 11 wild-type (WT) monkeys (Table S1) were used to construct the connectivity network using 94 parcellated brain regions (see Table S2 for a complete list of anatomical labels) for each dataset. The group lasso algorithm automatically and objectively identified 9 core regions out of 94 nodes of the monkey brain, which are the left central temporal cortex (CTC), right superior temporal cortex (STG), right dorsolateral prefrontal cortex (dlPFC), right primary somatosensory cortex (S1) and right primary motor cortex (M1), left anterior cingulate cortex (ACC), right centrolateral prefrontal cortex (clPFC), left superior parietal cortex (SPL), right ventrolateral prefrontal cortex (vlPFC) (Fig. 2A). To assess the biological significance of these core regions, a two sample *t*-test of node strength, i.e. the sum of functional connnectivities of all edges connecting this node was performed between TG and WT groups. Relative to WT group, the TG group exhibited significantly increased node strength in right vlPFC (*p* = 0.004*, * means FDR corrected for multiple comparisons thereinafter), left ACC (*p* = 0.008*), right clPFC (*p* = 0.001*) and right dlPFC (*p* = 0.004*) (Fig. 2B). In contrast, the right S1 (*p* = 0.002*) and right M1 (*p* = 0.025) showed decreased node strength in TG group (Fig. 2B). In addition, the correlation analyses between node strength and phenotypic behaviors observed in transgenic monkeys, measured by time in repetitive circular routing and locomotion (17, 28), revealed a significant negative relationship between the right vlPFC and circular routing (*r* = −0.932, *p* = 0.021) and between left STG and locomotion (*r* = −0.922, *p* = 0.026) (Fig. S1). Conversely, the node strength of left ACC was positively associated with relative circular routing in TG group (*r* = 0.910, *p* = 0.032) (Fig. S1).

**Fig. 2.**
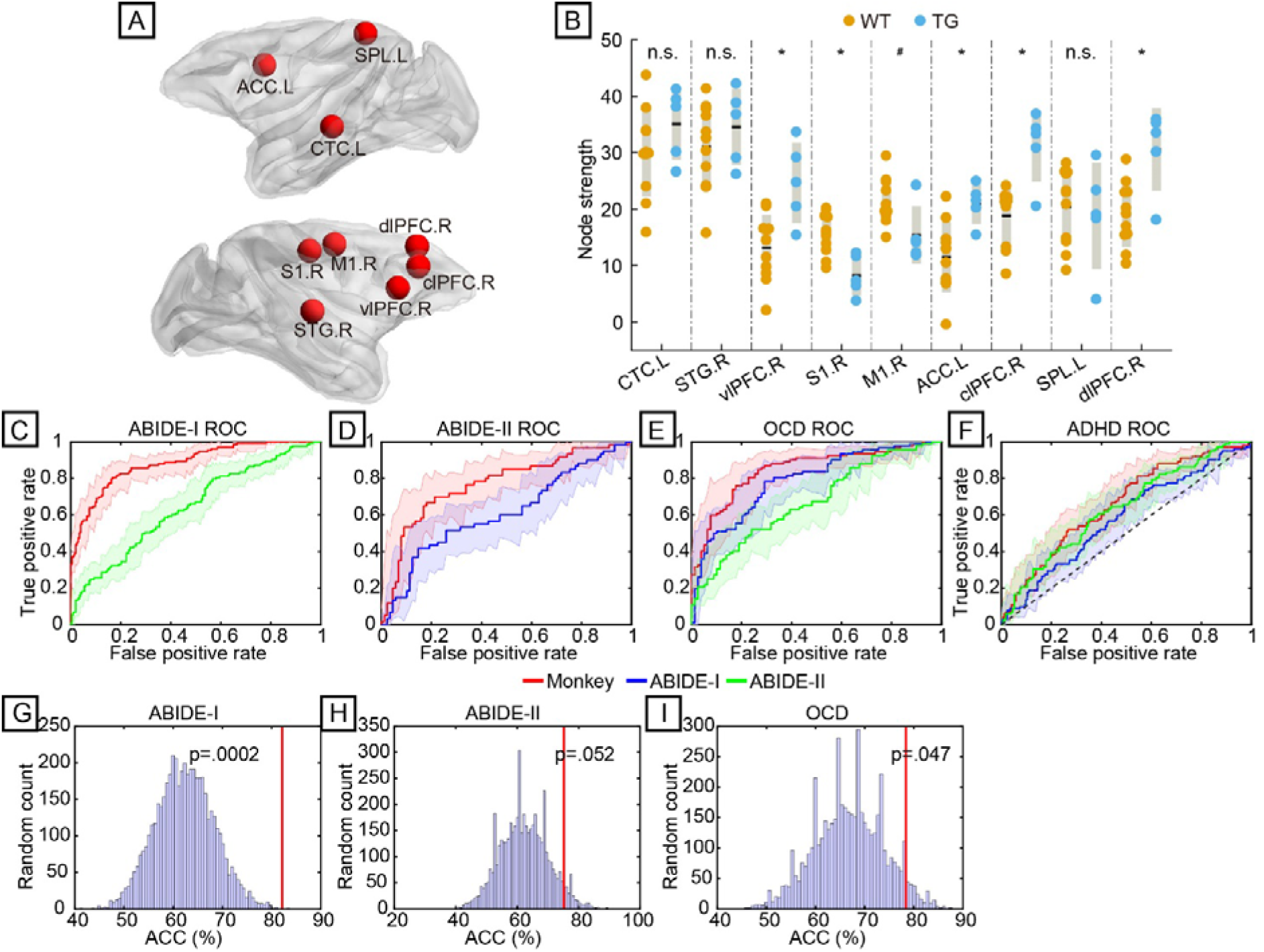
Performance of diagnostic classification for ASD and OCD cohorts using the monkey-/human-based classifiers. (A) Spatial distribution of 9 core regions identified from the monkey datasets. (B) Node strength of the 9 core regions in TG (*n* = 5, blue dots) compared with WT monkey (*n* = 11, yellow dots) using two sample *t*-test. Relative to WT group, the TG group exhibited significantly increased node strength in right vlPFC, left ACC, right clPFC and right dlPFC, whereas decreased node strength in right S1 and right M1. A black line denotes the group-averaged node strength and gray shading denotes standard deviation. * indicates *p* < 0.05, FDR corrected; # indicates *p* < 0.05, uncorrected; n.s., not significant. (C-F) Receiver operating characteristic (ROC) curves of the monkey-based classifier are plotted with those of the human-based classifier in ABIDE-I (C), ABIDE-II (D), OCD (E) and ADHD dataset (F), respectively. Red, blue and green lines denote ROC curves of the monkey-, the ABIDE-I and the ABIDE-II based classifier, respectively; shaded area indicates a 95% confidence interval. (G-I) Accuracy histograms of cross-validation by using randomly-generated core regions to construct the classifier in two ASD datasets and one OCD dataset. The vertical red lines indicate the classification accuracy achieved by the monkey-based classifier, which reaches a statistical confidence level at *p* = 0.0002 (G), *p* = 0.052 (H), and *p* = 0.047 (I).

### Highly accurate monkey-based classifier for ASD

Applying these 9 core regions to the human connectomics datasets, a functional-connection (FC), monkey-based classifier was constructed (there are a total of 801 FCs based on these 9 regions) (Fig. 1D). For diagnostic tests, 133 subjects with ASD and demographically matched 203 healthy controls (HC) were used from a total of 1112 subjects in ABIDE-I (8), and 60 subjects with ASD and 89 matched HC were used from 1114 subjects in ABIDE-II (22) after basic demographic and diagnostic screening. Using the standard lasso with a 10 x 10 nested cross-validation (29), we determined a subset of relevant, non-redundant edge features among the 801 FCs for use in classification. Note that the test set was never used for validation or feature selection (Fig. 1D; Fig. S2). A sparse logistic regression (30) in tandem with leave-one-participant-out cross validation was subsequently implemented to classify patients with ASD from HC in ABIDE-I cohort (Fig. 1E; Fig. S2), and was achieved with a cross-validated accuracy of 82.14% (Permutation test *p* = 0.001, Fig. S3; 95% confidence interval (CI), 77.53% to 86.00%), sensitivity of 79.70% (95% CI, 71.66% to 85.98%) and specificity of 83.74% (95% CI, 77.78% to 88.40%), corresponding to the area under the receiver operating characteristic (ROC) curve (AUC) of 0.884 (Fig. 2C; Table S3). In the independent ABIDE-II cohort, the monkey-based classifier also achieved remarkable performance with an accuracy of 75.17% (Permutation test *p* = 0.014, Fig. S3; 95% CI, 67.30% to 81.71%), sensitivity of 70.00% (95% CI, 56.63% to 80.80%), specificity of 78.65% (95% CI, 68.43% to 86.35%), and the AUC was 0.769 (Fig. 2D; Table S3).

We applied the same group lasso algorithm to one human ASD cohort to identify core regions, and used them to construct a classifier for classification in other ASD cohorts. Namely, the group lasso penalty was applied to identify core regions in one human ASD cohort, which were then used to construct the classifier for diagnostic classification in another human ASD cohort, and vice versa. In the ABIDE-II data set, we identified core regions as the left STG, left dorsal part of anterior visual area (VACd), left Visual area 2 (V2), right M1, bilateral ACC, right clPFC, right vlPFC, and right globus pallidus (GP) (Fig. S4). The classifier based on these core regions achieved an accuracy of 61.31% (95% CI, 55.85% to 66.51%), sensitivity of 56.39% (95% CI, 47.53% to 64.88%), specificity of 64.53% (95% CI, 57.49% to 71.02%) and AUC of 0.644 in the ABIDE-I cohort (Fig. 2C; Table S3), which is significantly lower than that of the monkey-based classifier (*p* < 0.001, McNemar’s test, Table S3) (31). By contrast, core regions identified in the ABIDE-I cohort were the left thalamus (THa), right primary visual area (V1), right V2 and right STG (Fig. S4). These ABIDE-I based classifier achieved an accuracy of 60.40% (95% CI, 52.04% to 68.21%) with a sensitivity of 53.33% (95% CI, 40.10% to 66.14%), specificity of 65.17% (95% CI, 54.26% to 74.76%) and AUC of 0.611 in the ABIDE-II cohort, which is significantly lower than that of the monkey-based classifier (*p* = 0.003, Fig. 2D; Table S3). Furthermore, we asked whether this monkey-based classifier would outperform random choice (by chance), i.e., randomly picking 9 out of 94 nodes as ‘core regions’. We repeated the above monkey-based classifier analysis through using a random selection of 9 regions and constructing the corresponding classifier to distinguish ASD from HC. This was repeated 5,000 times to assess the probability of attaining better accuracy than the actual monkey-based classifier. We observed that the performance of the monkey-based classifier was significantly better than chance for the ABIDE-I (*p* = 0.0002, Fig. 2G) and -II cohorts (*p* = 0.052, Fig. 2H). Note that there are some random selections showing higher accuracy than the monkey-based classifier (Fig. 2G,H). Upon closer examination, we found that a random selection of 9 regions outperforms the monkey-based one in one cohort whereas it always fails to achieve reasonable performance in other cohorts, indicating poor generalizability for the random choices of core regions (Fig. S5).

### Application of the monkey-based classifier to other human disorders

We next tested the generalizability of the monkey-based classifier in OCD and ADHD datasets. The OCD cohort contained 92 OCD patients and 79 healthy controls which were recruited through the local department of the OCD Clinic at Ruijin Hospital and part of this data set has recently been published elsewhere (23). The ADHD cohort that included 102 subjects with ADHD and 173 demographically matched HC was enrolled from a public ADHD-200 sample which provides a total of 776 subjects including 285 ADHD patients and 491 HC (24). In this OCD cohort, the classifier based on the same set of core regions from the monkey model achieve a remarkable performance with an accuracy of 78.36% (Permutation test *p* = 0.002, Fig. S3; 95% CI, 71.29% to 84.13%), sensitivity of 73.91% (95% CI, 63.53% to 82.26%), specificity of 83.54% (95% CI, 73.14% to 90.61%) and AUC of 0.848 (Fig. 2E), which outperformed other two human-based classifiers (ABIDE-I, *p* = 0.044; ABIDE-II, *p* = 0.000; Table S3), and the randomization classifiers (*p* = 0.047, Fig. 2I). However, in the ADHD cohort, the monkey-based classifier performed with an accuracy of 64.73% (95% CI, 58.73% to 70.31%), sensitivity = 51.96% (95% CI, 41.90% to 61.88%), specificity = 72.25% (95% CI, 64.85% to 78.65%) and AUC = 0.662, which yielded no significant difference relative to other human-based classifiers (Fig. 2F; Table S3). Notably core regions identified in the monkey model showed better generalizability than randomly-generated ones in ASD and OCD (Fig. S5).

### Characteristics of nodes and edges in the monkey-based classifier for ASD and OCD

We proceeded to quantitatively evaluate the relation between key elements (nodes and edges) of the monkey-based classifier and clinical symptoms in ASD and OCD. The SLR algorithm automatically identified 101, 74 and 64 FCs from the ABIDE-I, ABIDE-II and OCD data sets for reliable classification of patients and HCs, respectively (Fig. 3A-C). We characterized the spatial profiles of these FCs by calculating their intra-lobe and inter-lobe proportional distribution (Fig. 3D-F). Analysis of the spatial distribution of these classifiers shows that both ASD and OCD shared discriminant features of intra-lobe FCs mostly within the prefrontal lobe, whereas fronto-temporal and fronto-subcortical connections significantly epitomize their overall differences (Fig. 3G). In 48 of 60 participants with research-reliable Autism Diagnostic Observation Schedule (ADOS) scores from the ABIDE-II cohort, we found that node strengths of the right vlPFC (*r* = −0.333, *p* = 0.021), left CTC (*r* = −0.308, *p* = 0.033), right STG (*r* = −0.336, *p* = 0.020) and right S1 (*r* = −0.315, *p* = 0.029) exhibited significant correlations with the communication sub-scores of the ADOS (ADOS COMM) in patients (Fig. 4A; Table S4). Negative correlations between the node strengths of left ACC (*r* = −0.295, *p* = 0.042) and right dlPFC (*r* = −0.291, *p* = 0.045) and the ADOS COMM scores were also found in these ASD patients (Table S4). The node strength of the right vlPFC was found associated with Yale-Brown Obsessive Compulsive Scale (Y-BOCS) scores in OCD patients (*r* = −0.217, *p* = 0.038, Fig. 4A). In addition, we found a significant association between the HAM-A scores of OCD patients and the node strength of right SPL (*r* = −0.237, *p* = 0.023).

**Fig. 3.**
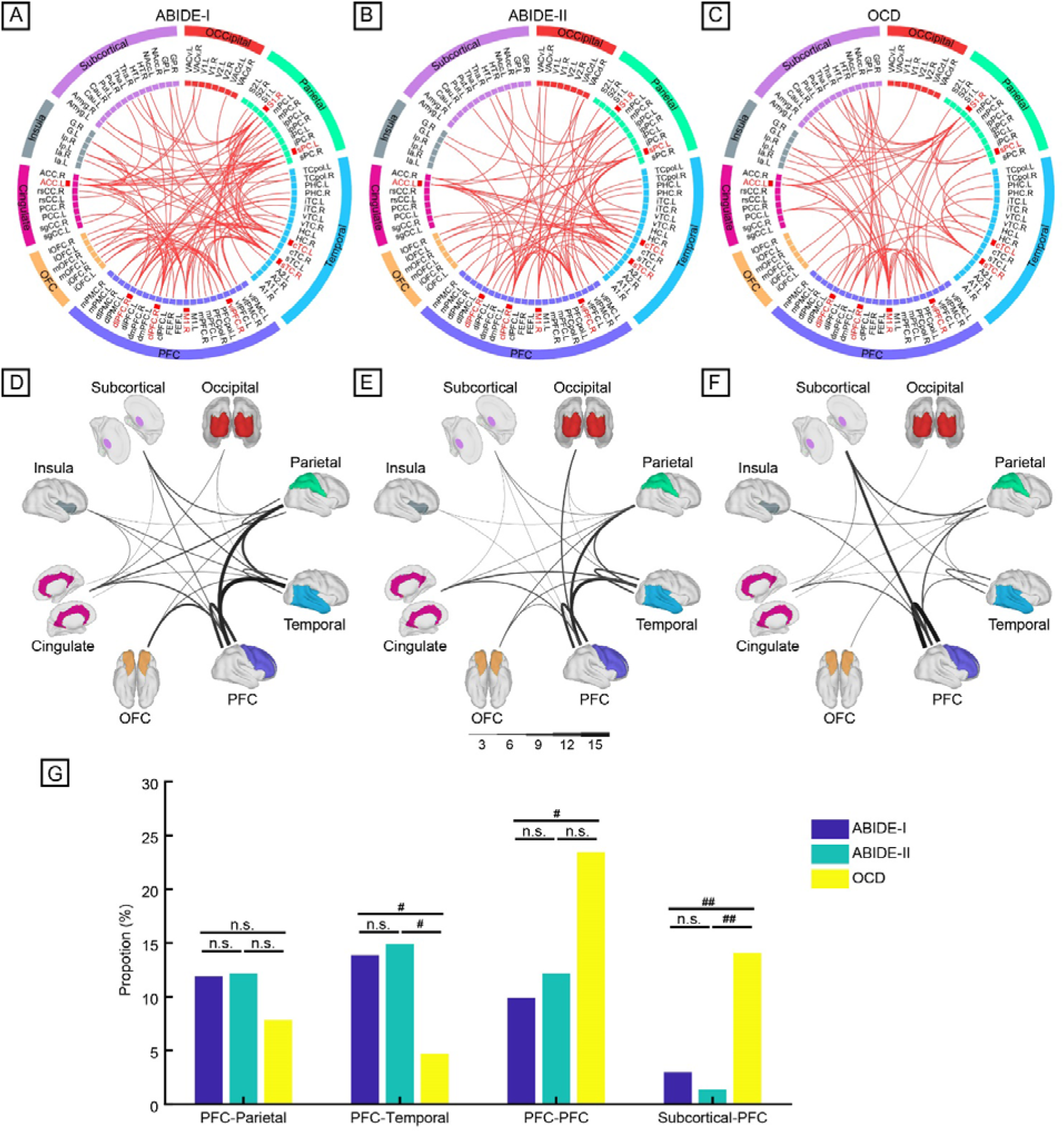
Characteristic of the identified functional connections for ASD and OCD. (A-C) The top row shows identified FCs and their terminal regions (nodes) for reliable distinguish ASD from HCs and OCD from HCs. The inner layer denotes 94 brain regions, and the interlayer highlights 9 core regions and the outer layer specifies the lobar division. (D-F) Middle row shows the sum of the number of intra- and inter-lobe connection based on the lobar division for ASD and OCD. (G) Bottom row shows the comparison of intra- and inter-lobe connection with relative higher FCs between ASD and OCD. # *p* < 0.05; ## *p* < 0.01uncorrected; n.s., not significant.

**Fig. 4.**
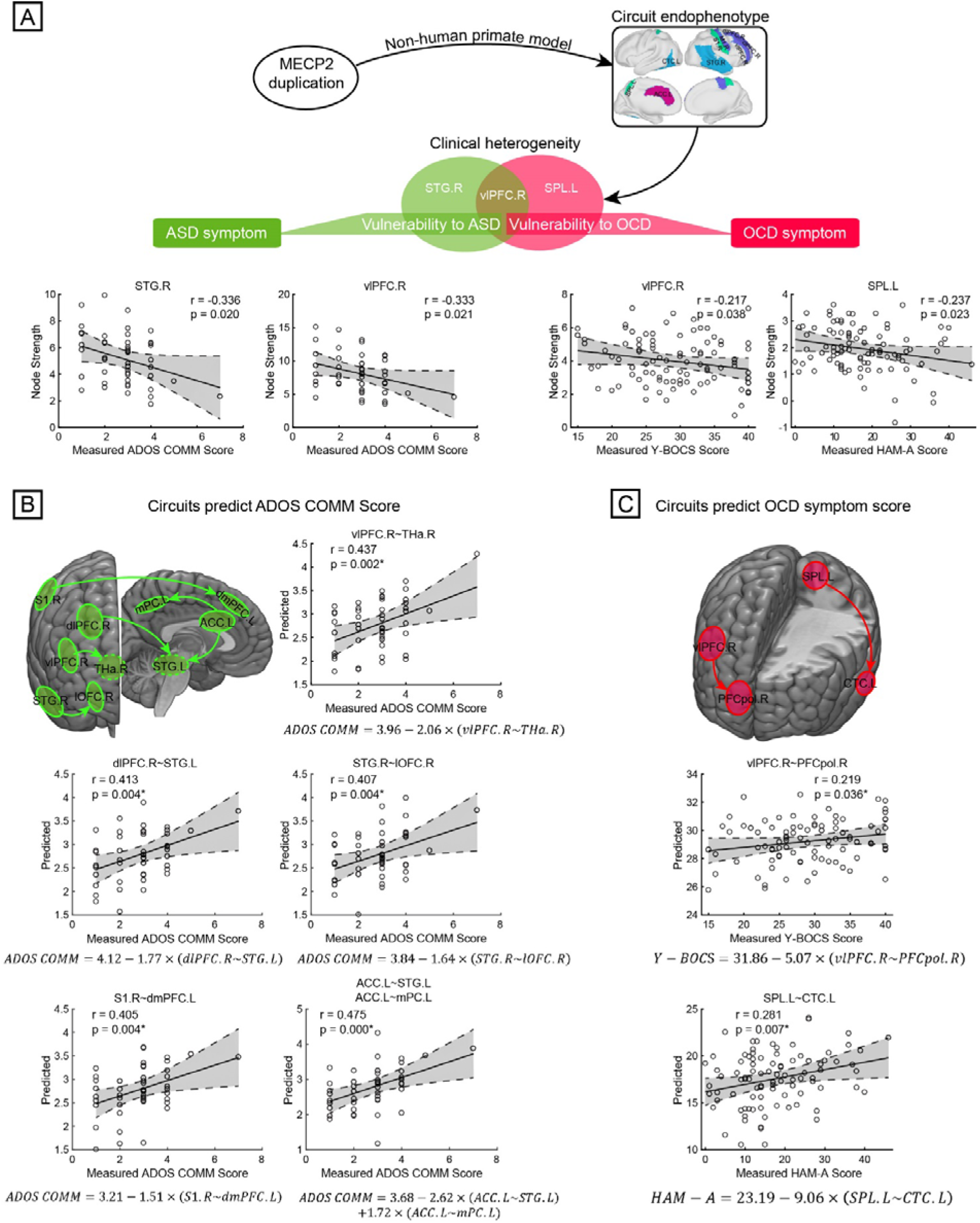
Genetically-linked neural circuits associated with distinct symptomatic domains in ASD and OCD. (A) Effects of single genetic mutation in animal models can be limited to some specific neural circuits (9 core regions) and associated behavioral manifestations in ASD and OCD. Node strengths of the right vlPFC and right STG exhibited significant correlations with the ADOS COMM scores of ASD patients, meanwhile the right vlPFC was found associated with Y-BOCS scores in OCD patients. And the right SPL was correlated with their HAM-A scores. (B) Predictions of ADOS COMM score based on linear models are shown at the bottom of each plot for core regions including right vlPFC, right dlPFC, right STG, right S1, and left ACC. The connection between the right vlPFC and right thalamus achieved a correlation of *r* = 0.437 (*p* = 0.002*) between predicted and measured ADOS COMM score in ABIDE-II patients. Similarly, the measured ADOS COMM score was predicted by the connections with right dlPFC and left STG (*r* = 0.413, *p* = 0.004*), right STG and right orbitolateral prefrontal cortex (*r* = 0.407, *p* = 0.004*), right S1 and left dorsomedial prefrontal cortex (*r* = 0.405, *p* = 0.004*), and the connections between left ACC and both left medial parietal cortex and left STG (*r* = 0.475, *p* < 0.001*), respectively. (C) Predictions of Y-BOCS and HAM-A scores in OCD based on linear model are shown at the bottom of plot for right vlPFC and left SPL. The connection between right vlPFC and right prefrontal polar cortex was sufficiently strong to predict the actual Y-BOCS scores of patients (*r* = 0.219, *p* = 0.036*). The actual HAM-A scores of these OCD patients were significantly predicted by the FC between left SPL and left CTC (*r* = 0.261, *p* = 0.007*). * indicates *p* < .05, FDR corrected. Shading indicates a 95% confidence interval.

For those regions exhibiting significant correlations with symptom severity scores, we performed a stepwise linear regression to predict symptom severity by modeling the relationship between the dependent variable (clinical symptom measurements) and the independent variables (FCs connected to a specific core region), which allowed determination of whether individual nodes and edges of the classifiers were commonly shared or uniquely linked to symptomatic domains of ASD and OCD. For instance, the FCs connecting to one core region such as the right vlPFC among all identified FCs were used to predict the ADOS COMM scores of ASD patients. Pearson’s correlation coefficient was used to evaluate the performance of the linear model in predicting symptom severity. The linear model based on the connection of right vlPFC-right THa achieved a correlation of *r* = 0.437 (*p* = 0.002*) between predicted and measured ADOS COMM scores in patients with ASD in the ABIDE-II group (*r* = 0.437, *p* = 0.002*, Fig. 4B; Table S5). The linear model is expressed as follows:

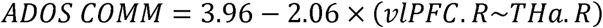

In fact, the ADOS COMM score was significantly predicted by connections between right dlPFC and left STG, right STG and right orbitolateral prefrontal cortex (lOFC), right S1 and left dorsomedial prefrontal cortex (dmPFC), between left ACC and both left medial parietal cortex (mPC) and left STG (Fig. 4B). By contrast, FC between right vlPFC and right prefrontal polar cortex (PFCpol) was predictive of the measured Y-BOCS scores of OCD patients (*r* = 0.219, *p* = 0.036*, Fig. 4C; Table S5):

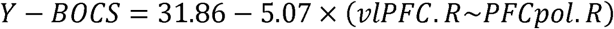

Of all left SPL-linked connections, the predicted HAM-A scores based on the FC between left SPL and left CTC was significantly correlated with the actual HAM-A scores of these OCD patients (*r* = 0.281, *p* = 0.007*, Fig. 4C; Table S5).

## Discussion

The superior predictive performance of the monkey-based model speaks to the value of investigating biologically homogeneous samples, particularly in the absence of sufficient real-world data available for deconstructing heterogeneity inherited in the clinical cohorts (given the overwhelming dimensions in most neuroimaging settings) (32). We have demonstrated that this particular set of core regions, although identified in a relatively small number of transgenic monkeys, is sufficiently powerful to set the stage for cross-species mapping and prioritization of features for reducing clinical heterogeneity. This is probably because predisposition of a single genetic manipulation in animal models can be highly susceptible to specific neural circuits/pathways, revealing the extent of a tangible circuit endophenotype and its manifestation at the behavioral domain (33). Fortunately, we managed to unravel a circuit-level path to successfully map key nodes in the pathological circuits of transgenic monkeys to the homologs in various human conditions, strongly suggesting an evolutionarily conserved mechanism of core regions in the functional specialization of large-scale brain networks.

Importantly, the present machine learning framework enables the search for some straightforward explanations for how a neural circuit allows superior classification for different human cohorts, and thus to gain illuminating insights into the mechanisms by which these features were linked to the etiology of ASD and OCD. This is illustrated by the fact that the core regions, mostly distributed in frontal and temporal lobes including right vlPFC right STG, were correlated with social communicative deficits (9, 21, 34). The right vlPFC and left ACC play an important role in the “stop circuit” (35, 36), heavily implicated in relation to restricted, repetitive behaviors and interests in ASD (37). Moreover, the FC between left SPL and left CTC predicted the severity of anxiety, and the relation between anxiety and attention in OCD (38). Additionally, our findings highlight the crucial role of two core regions S1 and M1, indicating their attributes to motor impairment in ASD (39) and impaired social-communicative skill development (39, 40), as deficits in S1 may affect the ability to master motor skills (41). Since sensory and motor disturbances have been hypothesized as some of the earliest signs of abnormality in children with ASD (40), future studies characterizing the trajectory of sensorimotor disturbance and the “downstream” effects onto social and communicative interaction in later development would be invaluable for addressing the mechanistic basis of symptomatic domains in autistic children.

From a circuit perspective, these core regions may simultaneously function as critical hubs to form differential patterns of functional coupling with the rest of the brain that underlie distinct clinical-symptom profiles in a variety of diseases. Upon close inspection, ASD-specific features exhibit apparently marked distribution of fronto-temporal and fronto-parietal connections, prominently involved in cognitive control and social communication (42). By contrast, dysconnectivity of OCD shows a significant bias towards the fronto-subcortical pathways, largely involved in response inhibition and cognitive flexibility (9, 35). Furthermore, a dissection of two distinct vlPFC-centered circuits from the monkey-based classifier has revealed dual neuropathology-dependent roles for the vlPFC which not only contributes to the communication domain of ASD but also to the compulsivity of OCD, thus emphasizing both unitary and diverse features contributing to the aetiological overlap between these disorders (14, 35).

This is to our knowledge the first study to propose this cross-species machine learning method for defining circuit endophenotypes and parsing heterogeneity and complexity across mental disorders, so caution is warranted. Replication to validate the present predictive algorithm with data from other populations and settings will be critical to extend the current use of genetically-engineered animal models to broader human application scenarios. Intriguingly, the combination of the 9 core regions pre-identified in transgenic monkeys effectively tease out relevant functional connections to achieve desirable performance compared to randomly-generated combinations (Fig. 2G-I), which calls for future biological validation regarding the role of the 9 core regions combination in the circuit pathology underlying both ASD and OCD and their potential utility in other disorders such as schizophrenia and addiction.

In summary, despite the heterogeneous genetic architecture of ASD and related disorders, the present translational strategy coupled with machine learning algorithms attests to the construct validity and feasibility of using genetically modified non-human primates to study human brain diseases, with important implication for parsing biologically-defined circuit endophenotypes that transcend conventional diagnostic boundaries in mental disorders. Nevertheless, the proposed interspecies framework can serve as a general strategy for bridging gaps between diagnostic categories of complex brain diseases, genetic and circuit mechanisms, thereby facilitating the development of targeted therapies based on symptom-specific mechanisms.

## Supplementary Materials

Materials and Methods

Supplementary References

Fig S1. Correlation between node strength of core regions and behavior abnormalities in TG group

Fig. S2. Illustration of nested 10×10 feature selection and leave-one-out cross-validation

Fig. S3. Null distribution of classification accuracy in the permutation test

Fig. S4. Identified core regions using ABIDE-I and ABIDE-II datasets

Fig. S5. Plots depicting the poor generalizability of random selections of 9 core regions in different cohorts

Table S1. Characteristics of all TG and WT monkeys

Table S2. Cortical and subcortical parcellation and abbreviations

Table S3. Classification performance of classifiers based on different sets of core regions

Table S4. Correlations between node strength and symptom severity in ASD and OCD cohorts

Table S5. Prediction of symptom severity using functional connections identified in the classifier

Table S6. Characteristics of human ABIDE-I cohorts

Table S7. Characteristics of human ABIDE-II cohorts

Table S8. Characteristics of human OCD cohorts

Table S9. Characteristics of human ADHD cohorts

Table S10. Imaging protocols for resting-state fMRI used in the present study

## Acknowledgements

We would like to thank Hu Zhang, Zhiwei Wang, Jinqiang Peng and Wenwen Yu for their assistance to the monkey data acquisition, thank Zhiwei Wang and Xiaoyu Chen for their assistance to the data preprocessing, and also thank Drs. Muming Poo, Ravi Menon, Valerie Voon for their stimulating discussions and suggestions during the preparation of this study.

## Author contributions

Z.W., R.H. and T.N.T conceived the study, T.W.R. and T.N.T. provided ongoing guidance on the design. R.H. and T.N.T. oversaw the data analysis using machine learning. T.W.R. and Z.W. oversaw the cross-species biological modeling and data interpretation. Y.F.Z., J.L., J.Z.W., with the help of Z.W. and R.H., analyzed the animal data. Y.F.Z., with the help of J.L., J.Z.W., X.X. and Z.W., analyzed the human data. Z.W. and T.W.R. wrote the manuscript with input from all authors. All authors read and approved the manuscript.

