## Supplemental Materials for "Cross-species machine learning improves diagnostic classification of human psychiatric disorders"

**This PDF file includes:**

Materials and Methods

Supplementary References

Fig. S1 to S5

Table S1-S10

### Materials and Methods

#### Animal preparation

All experimental procedures for nonhuman primate research in this study were approved by the Institutional Animal Care and Use Committee in the Institute of Neuroscience and by the Biomedical Research Ethics Committee, Shanghai Institutes for Biological Sciences, Chinese Academy of Sciences, and conformed to National Institutes of Health guidelines for the humane care and use of laboratory animals.

Monkey dataset consisted of five *MECP2*-duplication transgenic (TG) monkeys (*Macaca fascicularis*, age 4.40 ± 0.29 years (mean ± SD), weight 3.26 ± 0.75 kg; 2 male, 3 female) and 11 wild-type (WT) monkeys (*Macaca fascicularis*, age 4.68 ± 0.46 years, weight 3.97 ± 1.36 kg; 4 male, 7 female). All 16 monkeys were prepared and maintained in a stable brain state for fMRI scans. The fMRI scanning process of all monkeys was conducted in a similar manner to our previous work (*1*, *2*). Before each scanning session, anesthesia of the animals was inducted with an intramuscular injection of ketamine (10 mg per kg) and atropine sulfate (0.05 mg per kg). After intubation, animals were ventilated with an MRI-compatible ventilator (CWE Inc., Weston, Wisconsin). Macaques were maintained with intermittent positive-pressure ventilation to ensure a constant respiration rate (25-35 breaths/min). Local anesthetic (5% lidocaine cream) was applied around the ears to block peripheral nerve stimulation. The monkeys were then placed in a custom-built MRI-compatible stereotaxic frame with their belly facing downward, and their heads were secured before being inserted into the center of AC88 bores.

In light of the anesthesiologist's instructions, anesthesia was maintained using the lowest possible concentration of isoflurane gas. Isoflurane was selected for the scans as resting-state networks have previously been demonstrated to be present while using this agent (*1*, *3-5*). The vital signs of animals including blood oxygenation, ECG, rectal temperature (Small Animal Instruments, Inc., Stony Brook, New York), respiration rate and end-tidal $CO_2$ (Smiths Medical ASD Inc., Dublin, Ohio) were continuously monitored throughout the duration of the experiment. Oxygen saturation was kept at over 95% and body temperature was kept constant using a hot water blanket (Gaymar Industries Inc., Orchard Park, New York). Lactated Ringer's solution was given with a maximum rate of 10 ml/kg/hour during the anesthesia process (*6*). Note that our intention was to equate the levels of physiological anesthesia across animals and not the level of anesthetic gas concentration. Slight individual differences in physiology meant that slight differences in anesthetic gas concentrations were needed to impose a similar level of anesthesia on different monkeys (*7*). Brain states were monitored by simultaneous MRI-compatible electroencephalograph (EEG) (Brain Products GmbH, Gilching, Germany) during data acquisition. Within the range of isoflurane levels used in the current study, consistent patterns of functional coupling between distant brain areas have been reported in prior monkey fMRI studies (*5*, *8*) and demonstrated in our work as well (*1*, *2*).

#### Human Participant

*Human ASD cohort.* We analyzed data from ABIDE-I/II repository (*9*, *10*) and ADHD cohort from ADHD-200 (*11*), two publicly available multisite datasets of resting-state functional imaging data, and one OCD cohort from our institutional database. The ABIDE initiative now includes two large-scale collections: ABIDE-I and ABIDE-II. Each collection was created through the aggregation of datasets independently collected across more than 17 international brain imaging laboratories ([9], [10]). In the present study, two cohorts of human ASD data were selected from ABIDE-I and ABIDE-II separately. We included all individuals with a Diagnostic and Statistical Manual of Mental Disorders, Fourth Edition, Text Revision (*DSM-IV-TR)* diagnosis of either autism, Asperger syndrome, or pervasive developmental disorder not otherwise specified (PDD-NOS), collectively referred to as the ASD group and demographics matched healthy control (HC) subjects. As with other ABIDE studies ([10], [12]), participant inclusion criteria were as follows: (1) right-handedness; (2) a full-scale IQ (FIQ) score higher than 80; (3) ASD individuals with known current medication status; (4) a mean framewise displacement (FD) ([13]) of less than 0.2mm and the percent of frames or volumes with displacement greater than 0.2mm < 50%; (5) individuals with known eye status at scan (open/closed); (6) data with anatomical images providing near full brain coverage and successful registration; and (7) sites with at least 5 participants per group and matched number of participants between two groups after applying these inclusion criteria. This yielded data for 336 individuals (ASD = 133, male/female = 118/15, mean±SD age = 17.25±7.76 years; HCs = 203, male/female = 167/36, mean±SD age = 16.66±6.33 years) from 10 sites for ABIDE-I and 149 individuals (ASD = 60, male/female = 56/4, mean±SD age = 11.95±4.33 years; HCs = 89, male/female = 62/27, mean±SD age = 11.40±3.59 years) from 4 sites for ABIDE-II. More details of ASD samples are specified in Table S6 and S7.

*Human OCD cohort.* Between April 2013 and September 2016, patients were recruited through local inpatient and outpatient departments at the OCD Clinics at Ruijin Hospital. All participants provided written informed consent for study participation after receiving a complete description of the protocols, which were approved by the Institutional Review Boards at Ruijin Hospital, Shanghai Jiao

Tong University and by the Biomedical Research Ethics Committee, Shanghai Institutes for Biological Sciences, and Chinese Academy of Sciences. All patients had received a primary diagnosis of OCD based on clinical evaluation with the Chinese translation of the Structured Clinical Interview for *DSM-IV-TR*, and were administered the Yale-Brown Obsessive Compulsive Scale (Y-BOCS) (*14*) and Hamilton Anxiety Scale (HAM-A) to assess OCD symptom severity. Exclusion criteria were applied as follows: (1) translation or rotation in any axis of head motion larger than 3mm or 3° during scanning; (2) any neurological disorders, psychosurgery, current or past substance abuse or dependence, pregnancy or any substantial physical illness such as brain tumor, brain injury, stroke, or epilepsy. The OCD dataset consisted of 171 individuals (OCD = 92, male/female = 55/37, mean±SD age = 30.47±9.30 years; HCs = 79, male/female = 51/28, mean±SD age = 30.80±7.77 years). More details of OCD samples are specified in Table S8.

*Human ADHD cohort.* Open-access, freely accessible Attention Deficit and Hyperactivity Disorder (ADHD-200 Sample) database (http://fcon_1000.projects.nitrc.org/indi/adhd200/) was used to obtain datasets with resting-state data of individuals with ADHD and healthy controls. In the present study, the inclusion criteria included: (1) right-handedness; (2) an FIQ score higher than 80; (3) ADHD individuals with known current medication status; (4) individuals with images accepted after quality control. Individuals with translation or rotation in any axis of head motion larger than 3mm or 3° during scanning were excluded after applying the above inclusion criteria. These criteria led to the inclusion of data from the Kennedy Krieger Institute (“KKI”), New York University Child Study Center (“NYU”), Oregon Health & Science University (“OHSU”) and Peking (“Peking”) study-sites. Note that, for sites that contributed several functional runs per participant (NYU, OHSU), we only used the first functional run in the present study. Sites with less than 5 participants per group and a mismatched number of participants between two groups after the above criteria applied were excluded. We selected ADHD participants with known current medication status and with images accepted after quality control. This yielded data for 275 individuals (ADHD = 102, male/female = 85/17, mean±SD = 11.57±2.23 years; HCs = 173, male/female = 99/74, mean±SD age= 10.96±1.81 years) for ADHD. More details of ADHD samples are specified in Table S9.

#### Monkey MRI data acquisition and preprocessing

MRI images of all monkeys were acquired at the Institute of Neuroscience on a 3T whole-body scanner (Trio; Siemens Healthcare, Erlangen, Germany) running with an enhanced gradient coil insert (AC88; 80 mT/m maximum gradient strength, 800 mT/m/s maximum slew rate). A custom-built 8-channel phased-array transceiver coil was used for animal imaging sessions. Whole-brain resting-state fMRI data were collected using a gradient-echo echo-planar sequence (repetition time [TR] = 2000 ms; echo time [TE] = 29 ms; flip angle = 77°; slices = 32; matrix = 64 × 64; field of view = 96 × 96 mm; 1.5 × 1.5 $mm^2$ in plane resolution; slice thickness = 2.5 mm; GRAPPA factor = 2). For each session, 5 to 10 runs were acquired and each run consisted of 200 functional volumes. A pair of gradient echo images (TE: 4.22 ms and 6.68 ms) with the same orientation and resolution as EPI images were acquired to generate a field map for distortion correction of EPI images. High-resolution T1-weighted anatomical images were acquired using a MPRAGE sequence (TR = 2500 ms; TE = 3.12 ms; inversion time = 1100 ms; flip angle = 9°; acquisition voxel size = 0.5 × 0.5 × 0.5 $mm^3$; 144 sagittal slices). Six whole-brain anatomical volumes were acquired and further averaged for better brain segmentation and 3D cortical reconstruction. In the present experimental design, we did not intentionally collect different numbers of runs from each subject. In practice, however, the actual scan time for each experiment varied with the physiological status of different animals on that day, as is the case in most animal fMRI studies (5, 8, 15). In the final analysis, we included a total of 45 runs from TG and 99 runs from WT monkeys. Details of run numbers and physiological parameters for each animal are listed in **Table S1**.

Functional images of monkey and human brains were preprocessed using exactly the same strategy, which included slice timing correction, motion correction, coregistration with individual T1-weighted image, normalization to corresponding standard space, reslicing and spatial smoothing, regression of nuisance signals, removal of linear drift and temporal filtering (0.01 - 0.1 Hz). Specifically, the preprocessing of the monkey data were done using the SPM 8.0 toolbox (http://www.fil.ion.ucl.ac.uk/spm) and the FMRIB Software Library toolbox (FSL; https://fsl.fmrib.ox.ac.uk/fsl). The first 10 volumes were discarded. The field map images of each participant were then applied to compensate for the geometric distortion of EPI images caused by magnetic field inhomogeneity using FSL FUGUE. After slice timing correction and motion correction, the corrected images were normalized to standard space of the monkey F99 atlas (http://sumsdb.wustl.edu/sums/macaque_more.do) using an optimum 12-parameter affine transformation and nonlinear deformations, and then resampled to 2-mm cubic voxels and spatially smoothed with a 4 mm full-width at half-maximum (FWHM) isotropic Gaussian kernel. Six head motion parameters, ventricle and white matter signals were removed from the smoothed volumes using linear regression. Linear drift of the volumes was removed and a temporal filter was performed.

#### Human MRI data acquisition and preprocessing

Details available for ABIDE-I/II at http://fcon_1000.projects.nitrc.org/indi/abide/ and ADHD-200 at http://fcon_1000.projects.nitrc.org/indi/adhd200/. The imaging parameters of ASD and ADHD cohorts are listed in **Table S10**. OCD MRI data were collected using a Siemens Tim Trio 3T scanner (Erlangen, Germany). All participants underwent both functional and structural MRI scanning. Resting-state fMRI scans of the whole brain were acquired using a T2*-weighted EPI (echo planar imaging) sequence: repetition time (TR) = 3000 ms; echo time (TE) = 30 ms; flip angle, 90°; 47 axial slices; 3 mm slice thickness with no gap; and 300 volumes. High-resolution T1-weighted images used a magnetization prepared rapid gradient echo sequence: TR = 2300 ms; TE = 3 ms; inversion time, 1000 ms; flip angle, 9°;1×1×1 $mm^3$ spatial resolution. During the resting-state scan, participants were instructed to stay awake but relaxed, with their eyes closed, remain motionless, and refrain from thinking about anything in particular.

*ASD data preprocessing*. The preprocessing of ABIDE-I was performed by the Preprocessed Connectomes Project (PCP, http://preprocessed-connectomes-project.org/abide/index.html) using the Data Processing Assistant for Resting-State fMRI (DPARSF) Toolbox (*16*). Preprocessing steps included slice timing correction, motion correction, spatial normalization into MNI space, reslicing to 3 × 3 × 3 mm voxels and smoothing with a Gaussian kernel (FWHM = 6 mm). Friston-24 parameters of head motion, white matter and ventricle signals were regressed out, followed by linear drift correction and temporal filtering (0.01 - 0.1 Hz). For more details, readers are suggested to refer to the description in the PCP (http://preprocessed-connectomes-project.org/abide/dparsf.html). The same preprocessing streamline was applied to the ABIDE-II data set using DPARSF Toolbox.

*OCD data preprocessing.* Preprocessing for OCD data were the same as the ABIDE PCP pipeline except for minimal differences in some parameters, a smaller smoothing kernel (i.e. 2 mm), six head-motion parameters and global mean signals were regressed, and temporal filtering was applied (0.01 - 0.08HZ) for OCD.

*ADHD data preprocessing.* The preprocessing of ADHD data was performed by the Preprocessed Connectomes Project (PCP, http://preprocessed-connectomes-project.org/adhd200/) by the Athena team based on tools from the AFNI and FSL (*17*). The Athena pipeline involved removing the first four volumes to allow for magnetization to reach equilibrium, site-specific slice timing correction to the middle slice, deoblique dataset reorient into RPI orientation, motion correcting EPI volumes to the first (originally 5th) image of the time series, masking the dataset to exclude non-brain, averaging the volumes to create a mean image, co-registration of the mean EPI image to corresponding anatomic image, and writing fMRI data and mean image into template space at $4 \times 4 \times 4$ mm$^3$ resolution. Mean WM and CSF signals extracted using the masks calculated during s-MRI processing were included along with 6 head motion parameters and a third-order polynomial in voxelwise nuisance regression models to remove variation due to physiological noise, head motion, and scanner drifts from the time series. The resulting denoised time series were band-pass filter ($0.009 < f < 0.08$ Hz) voxel time courses to exclude frequencies not implicated in resting state functional connectivity and then spatially smoothed with a 6 mm FWHM Gaussian (*17*). For more details of the preprocessing pipeline see https://www.nitrc.org/plugins/mwiki/index.php/neurobureau:AthenaPipeline.

#### Monkey and human brain parcellation and network construction

The cortical organizations of both monkeys and humans were parcellated according to the Regional Map template (*18*, *19*). As the Regional Map parcellation does not include subcortical regions, subcortical parcellation for the two species was added on the basis of the INIA19 (*20*) and Freesurfer templates (*21*), respectively. This generated a whole brain template with a total of 94 regions of interest (ROIs) for both monkeys and humans (see **Table S2** for a complete list of all anatomical labels). Pearson's correlation coefficients between the mean time courses of any pair of regions were calculated to represent their functional connectivity, resulting in a 94 × 94 connectivity network matrix. Fisher's Z-transformation was then applied to the connectivity matrix which was subject to a covariates regression procedure for additional analysis. Both age (linear and quadratic), and gender were used as covariates for monkey and human data. Furthermore if the FIQ, site information, current medication status and eye status at scan were known, then they were used as additional covariates for human data.

#### Sparse linear regression model based on group lasso method

We adopted the sparse linear regression model based on group lasso method penalty (*22*) to identify a subset of core brain regions relevant to ASD from *MECP2* monkeys. The group lasso penalty is designed to eliminate a group of variables, treating all edges connected to one region (node) as a group, simultaneously. Put it in another way, we are more interested in selection of a subset of nodes that have non-zero coefficients in their connections to all the nodes in the graphs (or selection of rows in the adjacency matrix).

Let $\{(A^{(1)}, Y_1), \ldots, (A^{(M)}, Y_M)\}$ be the $M$ monkey sample of undirected adjacency matrices with $N$ nodes and with their class labels $y$. The adjacency matrices $A$ of each sample were reshaped into a one-dimensional vector and stacked, forming a feature matrix $B \in \mathbb{R}^{M \times P}$ ($M$ samples and $P$ features, $P = N \times (N-1)$). Let $y = (Y_1, \ldots, Y_M)$ denotes the $M$ dimensional response vector (e.g. the diagnostic label, that is $y = 1$ indicates TG and $y = 0$ indicates WT class). The sparse linear regression model based on group lasso penalty can then be formulated as

$$\min_{x \in \mathbb{R}^P} F(x) = \frac{1}{2}\left\| y - \sum_{n=1}^{N} [B]_n [x]_n \right\|_2^2 + \lambda \sum_{n=1}^{N} w_n \|[x]_n\|_2 \qquad (1)$$

where λ is a positive regularization parameter with 100 equally spaced points between 0.05 and 1. In this setting, the original feature matrix $B$ was partitioned into $N$ groups (treating all edges connected to one node as a group) $[B]_1, [B]_2, \ldots\ldots, [B]_N$ and $w_n$ denotes the weight for the $n$-th group. Note that, here we used screening rule proposed by Wang, Wonka and Ye (*23*) to solve group lasso efficiently. After solving the group lasso problem, we get the corresponding $N$ solution vector $[x]_1, [x]_2, \ldots\ldots, [x]_N$ and the dimension of $[x]_n$ is the same as the feature space in $[B]_n$. The relevant group features selected at each regularization parameter were combined by the union operation, to avoid the tweaking of such parameters.

#### Cross-species diagnostic classification for human individuals

The group lasso algorithm automatically and objectively identified 9 core regions from transgenic monkeys and were leveraged to pre-selecting all edges that connected to the regions from in human correlation matrix, comprising 801 unique FCs. To test the biological validity of those 9 core regions identified in the monkey cohort, two-sample *t*-tests with two tails were applied to evaluate statistical significance of group differences in the node strength of the core regions between TG and WT group.

The statistical significant level was set at $p < 0.05$, false discovery rate (FDR) corrected for multiple comparisons. We also performed regional Pearson's correlation analyses between node strength of the 9 core regions and behavior abnormalities, measured by time in repetitive circular routing and locomotion (*24*, *25*), to evaluate biological relevance of these regions in monkey cohort. Subsequently, the procedure for selecting relevant FCs from the pre-selected FCs (801 FCs), training a predictive model and assessing its generalization ability was carried out as a sequential process of 10×10 nested feature-selection (FS) and leave-one-out cross-validation. The whole data set was split into ten folds using a stratified approach, to keep an equal amount of (diagnosis, gender and site) combinations per fold. In each leave-one-out (LOO) cross-validation (CV) fold, all-but-one subjects were used to train an SLR classifier, while the remaining subject was used for evaluation. Before training SLR, it is necessary to select a subset of relevant, nonredundant connectivity features. Therefore, prior to LOOCV, nested FS was performed using lasso (*26*). Lasso identifies the latent relationships between FCs and the diagnostic label (**Fig. S2**).

The feature (FCs) selection procedure was similar to 10×10 nested cross-validation, with the difference being that the test set was never used for validation or feature (FCs) selection. In this way, lasso was trained on different subsamples of the data set, to increase the stability of the selected features. The 'test set' of the outer loop FS process was kept as a testing pool for LOOCV, whereas the ten folds of the inner loop FS were used to select features. Consequently, the LOOCV folds that belonged to the same testing pool of the outer loop FS shared the same reduced features. In the inner loop FS, the FS was completed using MATLAB's *Statistics and regression Toolbox* (Mathworks Inc.). FCs were selected using the default setting of the *lasso* function. The hyperparameter λ was estimated default by *lasso*. The features selected at each inner fold and λ were combined by the union operation, to include features that are important for any possible subsample (inner ten folds) of the training data set. Once the inner loop FS was executed, one sample was taken from the testing pool of the outer loop FS, and used as the test set of the LOOCV. The remaining samples were used to train SLR on the FCs retained during the inner loop FS (as illustrated in **Fig. S2**).

To predict the diagnostic label from the extracted features (optimal FCs, $z$), we employed logistic regression as the classifier. In logistic regression, a logistic function is used to define the probability of a participant belonging to the ASD class as

$$P(y = 1 \mid \hat{z}; w) = \frac{1}{1 + \exp(-w^T \hat{z})} \qquad (2)$$

where $y$ represents the diagnosis class label, that is $y = 1$ indicates ASD and $y = 0$ indicates HC class, respectively. $\hat{z} = [z^T, 1]^T \in \mathbb{R}^{k+1}$ is a feature vector with an augmented input. $w \in \mathbb{R}^{k+1}$ is the weight vector of the logistic function. SLR automatically selects the features related to the ASD label as input for the logistic function. In SLR, the probability distribution of the parameter vector is estimated using the hierarchical Bayesian estimation approach, in which the prior distribution of each element of the parameter vector is represented as a Gaussian distribution. Because of the automatic relevance determination property of the hierarchical Bayesian estimation method, some of the Gaussian distributions become sharply peaked at zero so that the irrelevant features are not used in the classification (*27*).

#### Model Validation, Comparison and Generalization

To test model robustness, a non-parametric permutation test was performed to determine whether the classification accuracy simply occurred by chance. The entire classification analysis was repeated using randomly shuffled group labels 5,000 times. This procedure estimated the null distribution of classification accuracy and the significant $p$ value was estimated by identifying the proportion of the total number for which the classification accuracy was greater than the observed one. Moreover, the reliability of the proposed cross-species translational framework was further tested under conditions where the 9 core regions were randomly selected from the 94 regions to evaluate whether the probability of getting accuracy values was significantly higher than chance level ($n$ = 5000 iterations). For model comparison, we compared the predictive accuracies of two classification models, i.e. monkey-based classifier and human-based classifier using McNemar's test (*28*). Specifically, the monkey-based classifier was constructed based on core regions identified from monkey data, the human-based classifier was constructed based on core regions identified from human ASD cohort. Finally, to test generalizability, the monkey-based classifier was applied to OCD and ADHD datasets.

#### Statistical analysis and association with symptom severity

To test the biological validity of the 9 core regions, two-sample $t$-test was applied to evaluate group difference in the node strength between TG and WT group in the monkey data. A significance threshold was set at $p < .05$, false discovery rate (FDR) corrected. We quantitatively evaluated the functional connections for reliable classification of patients and HCs for ASD and OCD. To facilitate data characterization and interpretation, we sorted connections based on lobar, i.e. prefrontal lobe, orbitofrontal lobe, temporal lobe, parietal lobe, occipital lobe, cingulate cortex, insula and subcortical area. We defined the intra-lobe connection as the absolute number of node-to-node FCs that belong to the same lobe, while the inter-lobe connection was defined by the absolute number of node-to-node FCs that belong to different lobes to evaluate spatial profile of discriminant functional connectivity. One-sided Fisher's exact test was applied to assess the shared and disorder-specific FCs between ASD and OCD. A significance threshold was set at one-tailed $p < .05$.

To investigate brain-behavior associations, we conducted Pearson's correlation analyses between node strength of the 9 core regions and behavior abnormalities in transgenic monkeys, measured by time spent in repetitive circular routing and locomotion (24, 25). In the human ASD and OCD cohorts, we first assessed the association between node strength of core regions and clinical symptom severity by Pearson's correlation. Significance threshold was set at $p < .05$. For core regions which showed significant correlations with symptom severity scores, the stepwise linear regression model was used to predict symptom severity by modeling the relationship between the dependent variable (symptom severity score) and independent variables (functional connections connected to a specific region). The Pearson correlation coefficient between the predicted values and its measured values was used to assess the performance of the regression model in predicting symptom severity. Significance threshold was set at $p < .05$, FDR corrected.

24. D. C. Cai, Z. W. Wang, T. T. Bo, S. Y. Yan, Y. L. Liu, Z. W. Liu, K. Zeljic, X. Y. Chen, Y. F. Zhan, X. Xu, Y. S. Du, Y. W. Wang, J. Cang, G. Z. Wang, J. Zhang, Q. Sun, Z. L. Qiu, S. J. Ge, Z. Ye, Z. Wang, MECP2 duplication causes aberrant GABA pathways, circuits and behaviors in transgenic monkeys: neural mappings to patients with autism. *Unpublished*, (Unpublished).

25. Z. Liu, X. Li, J. T. Zhang, Y. J. Cai, T. L. Cheng, C. Cheng, Y. Wang, C. C. Zhang, Y. H. Nie, Z. F. Chen, W. J. Bian, L. Zhang, J. Xiao, B. Lu, Y. F. Zhang, X. D. Zhang, X. Sang, J. J. Wu, X. Xu, Z. Q. Xiong, F. Zhang, X. Yu, N. Gong, W. H. Zhou, Q. Sun, Z. Qiu, Autism-like behaviours and germline transmission in transgenic monkeys overexpressing MeCP2. *Nature* **530**, 98-102 (2016).

26. R. Tibshirani, Regression shrinkage and selection via the Lasso. *J Roy Stat Soc B Met* **58**, 267-

288 (1996).

27. O. Yamashita, M. A. Sato, T. Yoshioka, F. Tong, Y. Kamitani, Sparse estimation automatically selects voxels relevant for the decoding of fMRI activity patterns. *Neuroimage* **42**, 1414-1429 (2008).

28. Q. McNemar, Note on the Sampling Error of the Difference between Correlated Proportions or Percentages. *Psychometrika* **12**, 153-157 (1947).

### Supplementary Figures

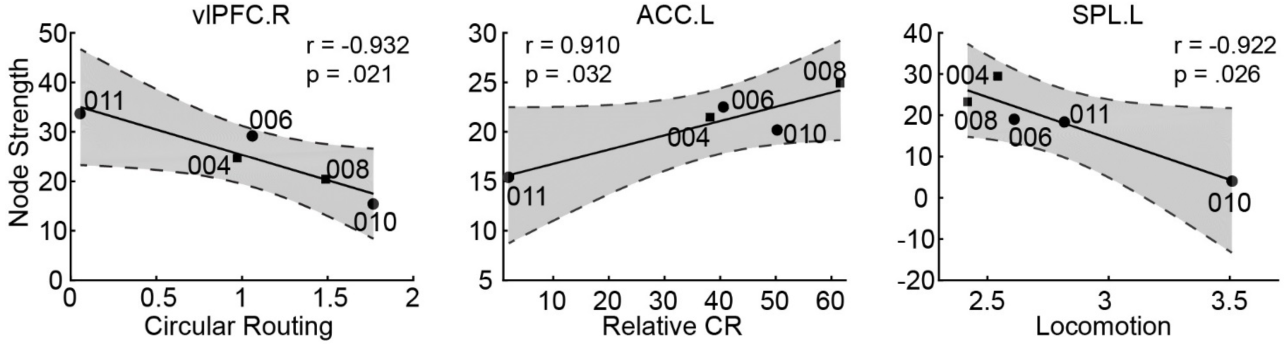


**Fig. S1. Correlation between node strength of core regions and behavior abnormalities in TG group.**

A circle shape denotes female and a square shape denotes male with labels indicating monkey ID. Gray zone indicating a 95% confidence interval. vlPFC.R, right ventrolateral prefrontal cortex; ACC.L, left anterior cingulate cortex; SPL.L, left superior parietal cortex; CR, circular routing.

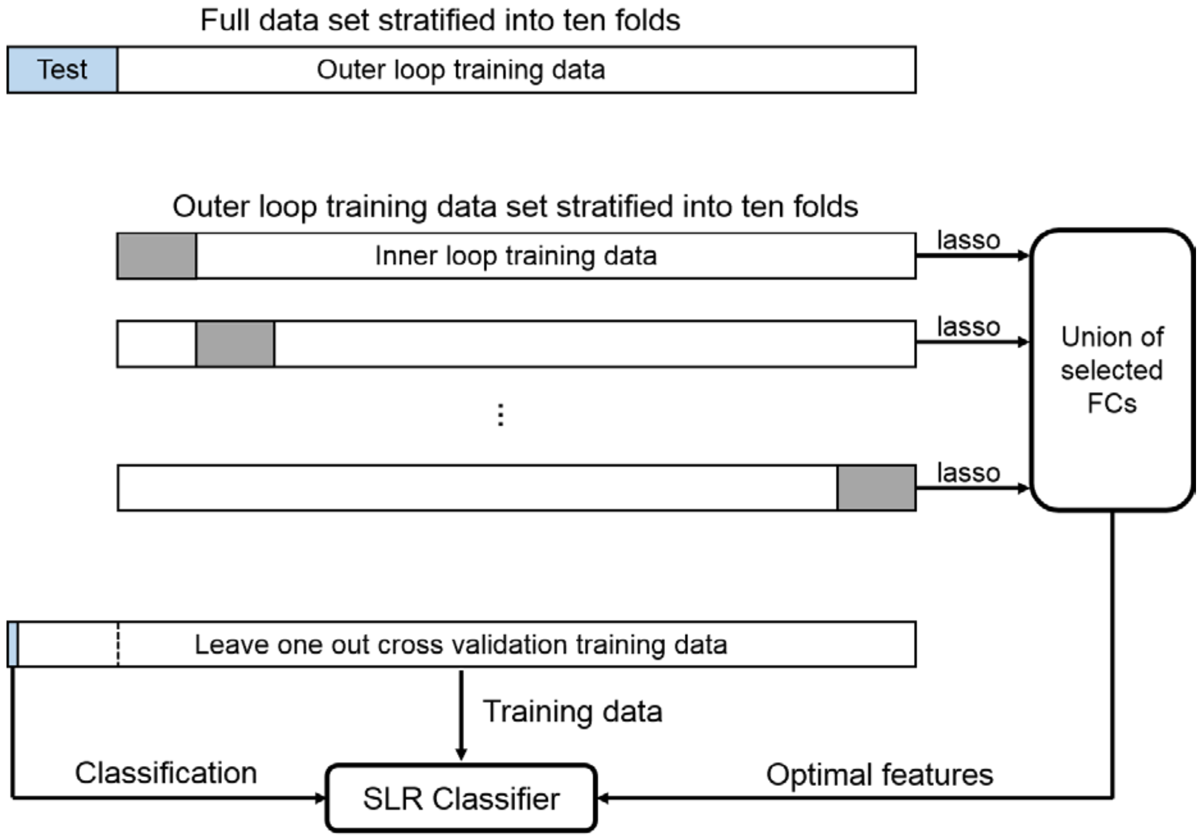


**Fig. S2. Illustration of nested 10×10 feature selection and leave-one-out cross-validation.**

I. Full data set was stratified into ten folds, one was first left out for testing, and the remaining 9 (outer loop training data) were used for constructing the optimal features.

II. In the inner loop, the outer loop training data was stratified into ten folds, one was first left out, and the remaining 9 (inner loop training data) were subjected to the lasso feature selection. The optimal features were union of the functional connections (FCs) selected throughout the inner loop.

III. In each iteration of the outer loop, one fold of the full sample is retained as a testing pool for leave-one-out cross-validation (LOOCV). One sample is taken from the testing pool and used as test set of LOOCV. The remaining samples are used to train SLR on optimal features retained during the inner loop. This procedure is repeated for every sample in the testing pool of the outer loop.

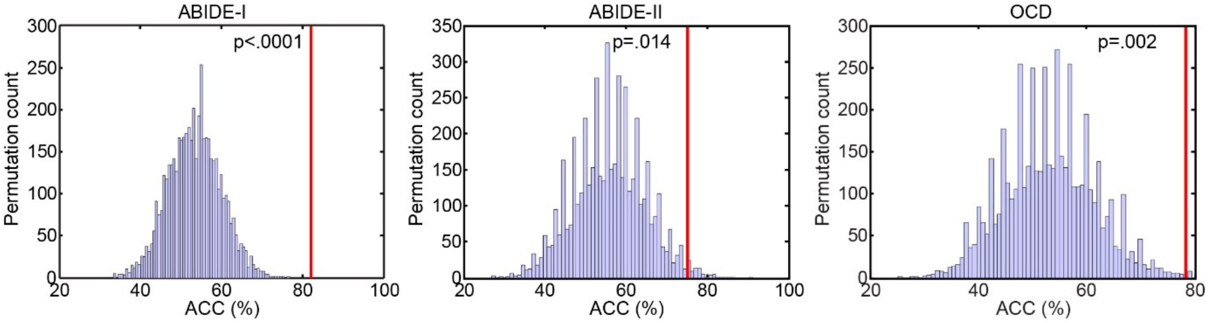


**Fig. S3. Null distribution of classification accuracy in the permutation test.**

The histograms of the permutation test (5,000 repetitions) for ASD (ABIDE-I and ABIDE-II) and OCD data. The purple bars represent the number of permutation counts at each level of accuracy. The vertical red lines denote the observed accuracies, corresponding to a p-value of $p$ = .000 for ABIDE-I, $p$ = .014 for ABIDE-II, and $p$ = .002 for OCD.

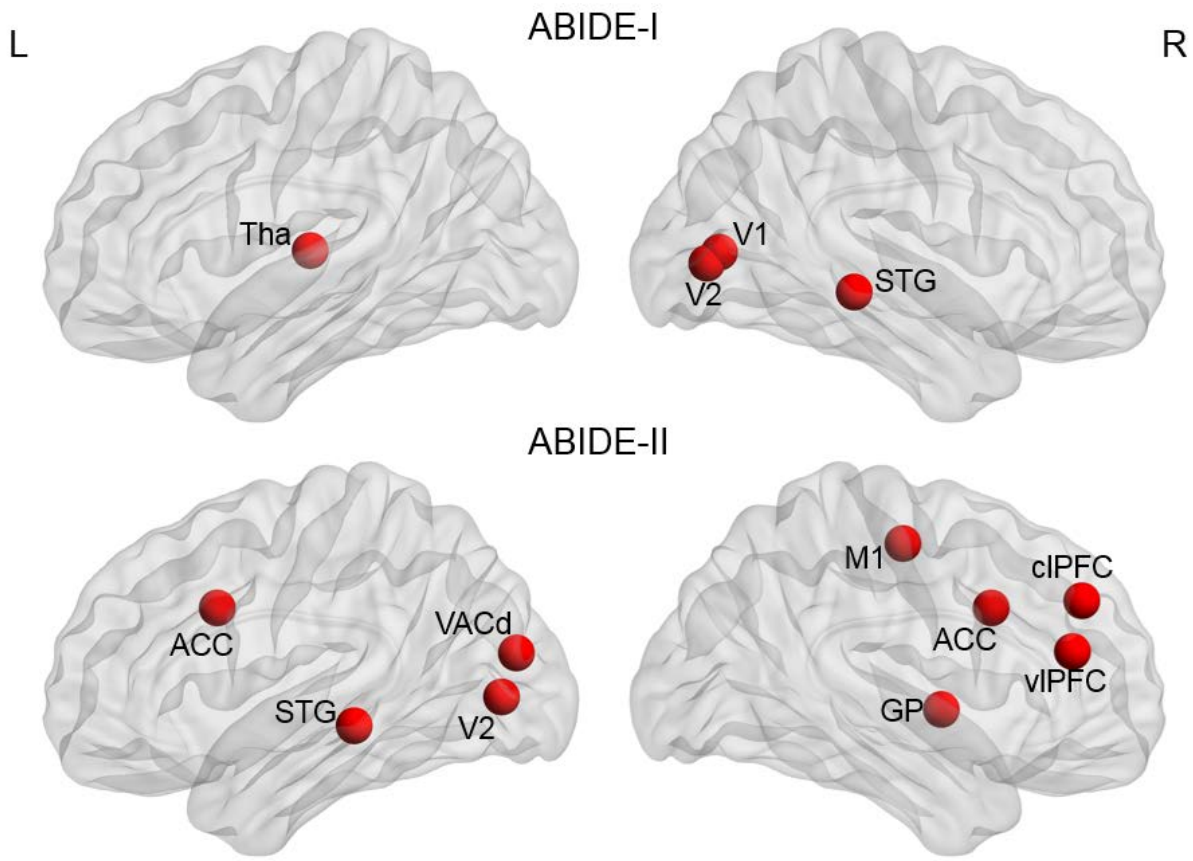


**Fig. S4. Identified core regions using ABIDE-I and ABIDE-II datasets.**

Core regions identified by group lasso in ABIDE-I and ABIDE-II cohorts. Tha, thalamus; V1, primary visual cortex; V2, secondary visual cortex; STG, superior temporal cortex; ACC, anterior cingulate cortex; VACd, ; M1, primary motor cotex; GP, globus pallidus; clPFC, centrolateral prefrontal cortex; vlPFC, ventrolateral prefrontal cortex.

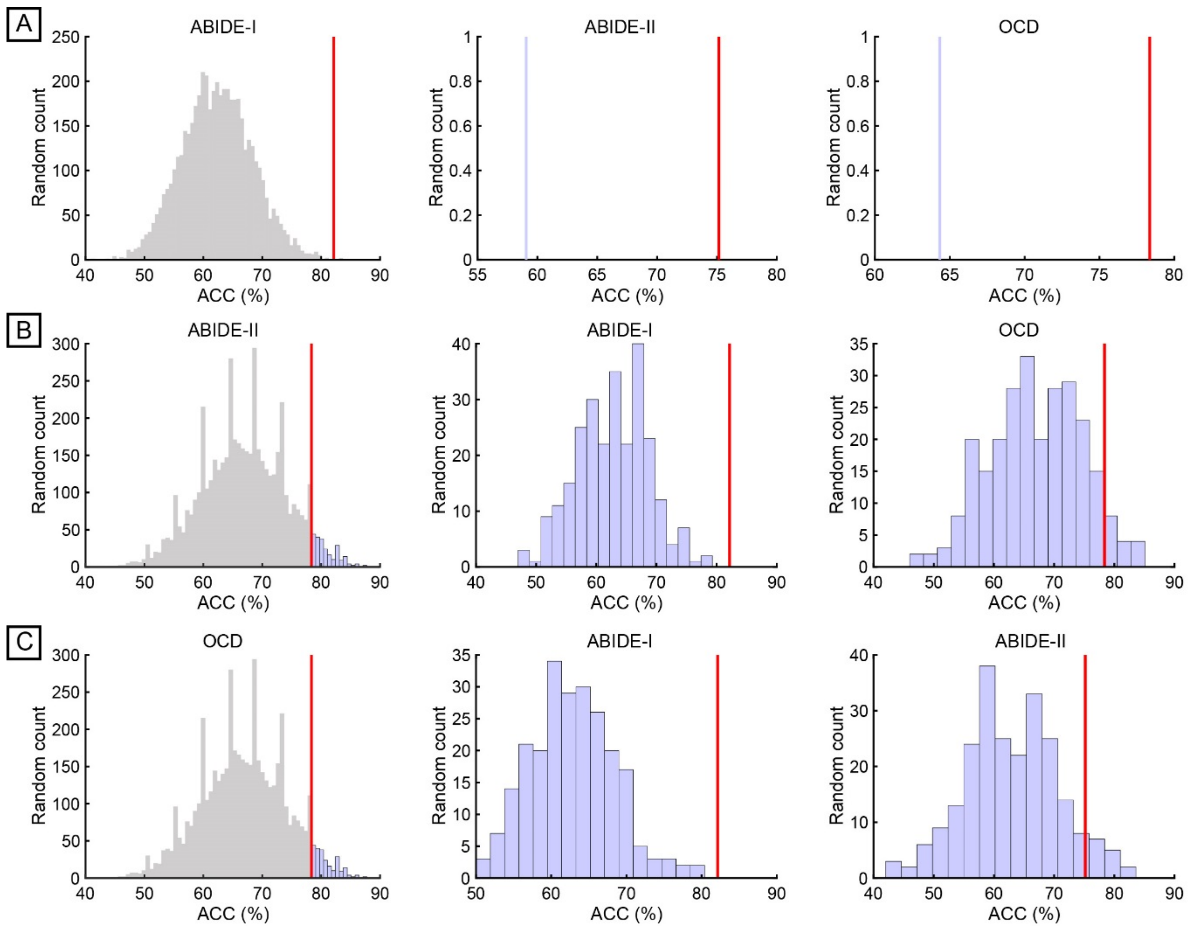


**Fig. S5. Plots depicting the poor generalizability of random selections of 9 core regions in different cohorts.**

Left column shows the accuracy histograms of 9 core regions that were randomly selected for 5,000 times in each dataset. Purple bars on the right side of the vertical red line denote cases with higher accuracy than the present monkey-based classifier. Middle and right columns show the accuracy of these cases compared to the monkey-based classifier in other two datasets.

### Supplementary Tables

**Table S1. Characteristics of all TG and WT monkeys.**

| | ID | Gender | Copy number | Weight (kg) | Age (year) | Heart rate (beat/min) | $EtCO_2$ (mmHg) | Temp. (°C) | Isoflurane (%) | Run |
|---|---|---|---|---|---|---|---|---|---|---|
| TG | TG04 | M | 1.0 | 4.3 | 4.9 | ~115 | ~29 | ~37.0 | 1.0 | 9 |
| | TG06 | F | 7.3 | 2.7 | 4.3 | ~103 | ~24 | ~35.6 | 1.0 | 8 |
| | TG08 | M | 2.9 | 3.8 | 4.2 | ~125 | ~29 | ~36.1 | 1.0 | 7 |
| | TG10 | F | 1.1 | 2.6 | 4.2 | ~94 | ~26 | ~35.8 | 1.0 | 8 |
| | TG11 | F | 1.9 | 2.9 | 4.4 | ~130 | ~27 | ~36.8 | 1.2 | 13 |
| | | 2M3F | | $3.26\pm0.75^{a}$ | $4.40\pm0.29^{a}$ | | | | | 45 |
| WT | WT030 | F | - | 3.5 | 4.8 | ~148 | ~27 | ~37.7 | 0.8 | 8 |
| | WT032 | M | - | 3.9 | 4.4 | ~120 | ~30 | ~37.7 | 1.0 | 10 |
| | WT034 | M | - | 3.5 | 4.4 | ~120 | ~30 | ~37.0 | 1.0 | 7 |
| | WT139 | M | - | 7 | 5.5 | ~148 | ~27 | ~37.2 | 1.2 | 10 |
| | WT278 | F | - | 3.4 | 4.5 | ~125 | ~28 | ~35.6 | 1.2-1.25 | 10 |
| | WT330 | F | - | 4.3 | 4.6 | ~125 | ~27 | ~35.9 | 1.2-1.3 | 10 |
| | WT358 | F | - | 3.2 | 4.2 | ~130 | ~28 | ~37.8 | 1.25-1.3 | 10 |
| | WT362 | F | - | 3 | 4.5 | ~130 | ~27 | ~36.1 | 1.2 | 10 |
| | WT463 | M | - | 6.1 | 5.1 | ~110 | ~27 | ~37.2 | 1.3-1.2 | 5 |
| | WT490 | F | - | 2.9 | 4.1 | ~136 | ~30 | ~37.1 | 1.1 | 9 |
| | WTTT | F | - | 2.9 | 5.4 | ~159 | ~27 | ~37.0 | 1.2 | 10 |
| | | 4M7F | | $3.97\pm1.36^{a}$ | $4.68\pm0.46^{a}$ | | | | | 99 |
| *p*-value | | $0.89^{b}$ | | | $0.233^{c}$ | | | | | |

Abbreviations. $EtCO_2$, end tidal of $CO_2$; Temp., rectal temperature; TG, transgenic; WT, wild-type.

a, mean ± SD; b, $\chi^2$ test; c, two sample student’s t-test with two tails.

**Table S2. Cortical and subcortical parcellation and abbreviations.**

| Lobes | Number | Hemisphere | Number | Hemisphere | Abbreviation | Full name |
|---|---|---|---|---|---|---|
| Occipital | 1 | L | 2 | R | V1 | Visual area 1 (primary visual cortex) |
| | 3 | L | 4 | R | V2 | Visual area 2 (secondary visual cortex) |
| | 5 | L | 6 | R | VACv | Anterior visual area, ventral part |
| | 7 | L | 8 | R | VACd | Anterior visual area, dorsal part |
| Parietal | 9 | L | 10 | R | S1 | Primary somatosensory cortex |
| | 11 | L | 12 | R | S2 | Secondary somatosensory cortex |
| | 13 | L | 14 | R | mPC | Medial parietal cortex |
| | 15 | L | 16 | R | IPS | Intraparietal cortex |
| | 17 | L | 18 | R | IPL | Inferior parietal cortex |
| | 19 | L | 20 | R | SPL | Superior parietal cortex |
| Temporal | 21 | L | 22 | R | A1 | Primary auditory cortex |
| | 23 | L | 24 | R | A2 | Secondary auditory cortex |
| | 25 | L | 26 | R | TCpol | Temporal polar cortex |
| | 27 | L | 28 | R | IT | Inferior temporal cortex |
| | 29 | L | 30 | R | VTC | Ventral temporal cortex |
| | 31 | L | 32 | R | CTC | Central temporal cortex |
| | 33 | L | 34 | R | CTC | Superior temporal cortex |
| | 35 | L | 36 | R | HC | Hippocampus |
| | 37 | L | 38 | R | PHC | Parahippocampal cortex |
| PFC | 39 | L | 40 | R | M1 | Primary motor cortex |
| | 41 | L | 42 | R | vlPMC | Ventrolateral premotor cortex |
| | 43 | L | 44 | R | dlPMC | Dorsolateral premotor cortex |
| | 45 | L | 46 | R | mPMC | Medial premotor cortex |
| | 47 | L | 48 | R | FEF | Frontal eye field |
| | 49 | L | 50 | R | vlPFC | Ventrolateral prefrontal cortex |
| | 51 | L | 52 | R | clPFC | Centrolateral prefrontal cortex |
| | 53 | L | 54 | R | dlPFC | Dorsolateral prefrontal cortex |
| | 55 | L | 56 | R | dmPFC | Dorsomedial prefrontal cortex |
| | 57 | L | 58 | R | mPFC | Medial prefrontal cortex |
| | 59 | L | 60 | R | PFCpol | Prefrontal polar cortex |
| OFC | 61 | L | 62 | R | iOFC | Orbitoinferior prefrontal cortex |
| | 63 | L | 64 | R | mOFC | Orbitomedial prefrontal cortex |
| | 65 | L | 66 | R | lOFC | Orbitolateral prefrontal cortex |
| Cingulate | 67 | L | 68 | R | sgACC | Subgenual cingulate cortex |
| | 69 | L | 70 | R | PCC | Posterior cingulate cortex |
| | 71 | L | 72 | R | rsCC | Retrosplenial cingulate cortex |
| | 73 | L | 74 | R | ACC | Anterior cingulate cortex |
| Insula | 75 | L | 76 | R | G | Gustatory cortex |
| | 77 | L | 78 | R | Ia | Anterior insula |
| | 79 | L | 80 | R | Ip | Posterior insula |
| Subcortical | 81 | L | 82 | R | Amyg | Amygdala |

| | | | | | |
|---|---|---|---|---|---|
| 83 | L | 84 | R | Cau | Caudate |
| 85 | L | 86 | R | Put | Putamen |
| 87 | L | 88 | R | Tha | Thalamus |
| 89 | L | 90 | R | HT | Hypothalamus |
| 91 | L | 92 | R | NAcc | Nucleus accumbens |
| 93 | L | 94 | R | GP | Globus pallidus |

**Table S3. Classification performance of classifiers based on different sets of core regions.**

| | Data set used to identify core regions | ACC (%) [95% CI] | Sensitivity (%) [95% CI] | Specificity (%) [95% CI] | AUC | *p* |
|---|---|---|---|---|---|---|
| ABIDE-I | Monkey | 82.14 [77.53%, 86.00%] | 79.70 [71.66%, 85.98%] | 83.74 [77.78%, 88.40%] | 0.884 | <.0001 |
| | ABIDE-II | 61.31 [55.85%, 66.51%] | 56.39 [47.53%, 64.88%] | 64.53 [57.49%, 71.02%] | 0.644 | |
| ABIDE-II | Monkey | 75.17 [67.30%, 81.71%] | 70.00 [56.63, 80.80%] | 78.65 [68.43%, 86.35%] | 0.769 | .003 |
| | ABIDE-I | 60.40 [52.04%, 68.21%] | 53.33 [40.10%, 66.14%] | 65.17 [54.26%, 74.76%] | 0.611 | |
| OCD | Monkey | 78.36 [71.29%, 84,13%] | 73.91 [63.53%, 82.26%] | 83.54 [73.14%, 90.61%] | 0.848 | -- |
| | ABIDE-I | 69.59 [62.02%, 76.26%] | 63.04 [52.29%, 72.69%] | 77.22 [66.15%, 85.59%] | 0.790 | .044[a] |
| | ABIDE-II | 60.23 [52.45%, 67.54%] | 57.61 [46.87%, 67.71%] | 63.29 [51.64%, 73.64%] | 0.674 | .000[b] |
| ADHD | Monkey | 64.73 [58.73%, 70.31%] | 51.96 [41.90%, 61.88%] | 72.25 [64.85%, 78.65%] | 0.662 | -- |
| | ABIDE-I | 58.18 [52.09%, 64.03%] | 49.02 [39.06%, 59.05%] | 63.58 [55.90%, 70.66%] | 0.582 | .067[a] |
| | ABIDE-II | 60.36 [54.29%, 66.14%] | 46.08 [36.26%, 56.20%] | 68.79 [61.24%, 75.49%] | 0.637 | .213[b] |

The comparison between different classifiers was conducted using McNemar's test. [a] denotes the monkey-based classifier in comparison with the ABIDE-I based classifier. [b] denotes the monkey-based classifier in comparison with the ABIDE-II based classifier.

ACC, accuracy; AUC, area under the receiver operating characteristic curve.

**Table S4. Correlations between node strength and symptom severity in ASD and OCD cohorts.**

| Core region | ABIDE-II | | | | OCD | | | |
|---|---|---|---|---|---|---|---|---|
| | ADOS TOTAL | | ADOS COMM | | Y-BOCS | | HAM-A | |
| | *r* | *p* | *r* | *p* | *r* | *p* | *r* | *p* |
| CTC.L | -0.307 | .034 | -0.308 | .033 | -0.082 | .44 | -0.199 | .058 |
| STG.R | -0.278 | .056 | -0.336 | .020 | -0.050 | .636 | -0.209 | .046 |
| vlPFC.R | -0.282 | .052 | -0.333 | .021 | -0.217 | .038 | -0.209 | .046 |
| S1.R | -0.294 | .043 | -0.315 | .029 | -0.132 | .208 | -0.124 | .237 |
| M1.R | -0.284 | .050 | -0.264 | .070 | -0.114 | .277 | -0.166 | .113 |
| ACC.L | -0.219 | .134 | -0.295 | .042 | -0.085 | .423 | -0.091 | .390 |
| clPFC.R | -0.138 | .351 | -0.242 | .098 | -0.134 | .204 | -0.165 | .117 |
| SPL.L | -0.170 | .249 | -0.209 | .153 | -0.158 | .132 | -0.237 | .023 |
| dlPFC.R | -0.226 | .123 | -0.291 | .045 | -0.182 | .083 | -0.091 | .387 |

CTC.L, left central temporal cortex; STG.R, right superior temporal cortex; vlPFC.R, right ventrolateral prefrontal cortex; S1.R, right primary somatosensory cortex; M1.R, right primary motor cortex; ACC.L, left anterior cingulate cortex; clPFC.R, right centrolateral prefrontal cortex; SPL, left superior parietal cortex; dlPFC.R, right dorsolateral prefrontal cortex. ADOS, Autism Diagnostic Observation Schedule; ADOS COMM, communication total sub-score of the classic ADOS; Y-BOCS, Yale-Brown Obsessive Compulsive Scale; HAM-A, Hamilton Anxiety Rating Scale.

**Table S5. Prediction of symptom severity using functional connections identified in the classifier.**

| Data | Core region | Predicted model | *r* | *p* |
|---|---|---|---|---|
| ABIDE-II | STG.R | $ADOS\ COMM = 3.84 - 1.64 \times (STG.R{\sim}lOFC.R)$ | 0.407 | .004* |
| | vlPFC.R | $ADOS\ COMM = 3.96 - 2.06 \times (vlPFC.R{\sim}THa.R)$ | 0.437 | .002* |
| | S1.R | $ADOS\ COMM = 3.21 - 1.51 \times (S1.R{\sim}dmPFC.L)$ | 0.405 | .004* |
| | ACC.L | $ADOS\ COMM = 3.68 - 2.62 \times (ACC.L{\sim}STG.L)$ $+1.72 \times (ACC.L{\sim}mPC.L)$ | 0.475 | <.0001* |
| | dlPFC.R | $ADOS\ COMM = 4.12 - 1.77 \times (dlPFC.R{\sim}STG.L)$ | 0.413 | .004* |
| OCD | vlPFC.R | $Y - BOCS = 31.86 - 5.07 \times (vlPFC.R{\sim}PFCpol.R)$ | 0.219 | .036* |
| | SPL.L | $HAM - A = 23.19 - 9.06 \times (SPL.L{\sim}CTC.L)$ | 0.281 | .007* |

CTC.L, left central temporal cortex; STG.R, right superior temporal cortex; vlPFC.R, right ventrolateral prefrontal cortex; S1.R, right primary somatosensory cortex; M1.R, right primary motor cortex; ACC.L, left anterior cingulate cortex; clPFC.R, right centrolateral prefrontal cortex; SPL, left superior parietal cortex; dlPFC.R, right dorsolateral prefrontal cortex; FEF.R, right frontal eye field; mPC.L, left Medial parietal cortex; iOFC, right orbitoinferior prefrontal cortex; THa.R, right thalamus; PFCpol.R, right prefrontal pole. * indicates $p < 0.05$, FDR corrected.

**Table S6. Characteristics of human ABIDE-I cohorts.**

| Site | Number | | Autism/Asperger/PDD-NOS | Age, mean±SD [Range] | | | Gender (M/F) | | | FIQ, mean±SD [Range] | | |
|---|---|---|---|---|---|---|---|---|---|---|---|---|
| | HC | ASD | | HC | ASD | $p^a$ | HC | ASD | $p^b$ | HC | ASD | $p^a$ |
| KKI | 21 | 5 | 1/4/0 | 10.10±1.15 [8.39 12.77] | 9.87±1.52) [8.09 11.37] | .701 | 14/7 | 4/1 | .562 | 113.14±9.54 [98.00 125.00] | 98.60±15.29 [84.00 120.00] | .012 |
| LEUVEN_1 | 14 | 13 | 13/0/0 | 23.36±3.00 [18.00 29.00] | 21.77±4.27 [18.00 32.00] | .271 | 14/0 | 13/0 | -- | 113.43±12.15 [98.00 146.00] | 108.00±12.44 [89.00 128.00] | .262 |
| MAX_MUN | 24 | 12 | 0/12/0 | 25.92±8.32 [7.00 46.00] | 32.08±14.46 [11.00 58.00] | .112 | 23/1 | 10/2 | .201 | 112.13±9.78 [95.00 129.00] | 114.67±12.62 [93.00 133.00] | .509 |
| PITT | 20 | 18 | 18/0/0 | 18.87±6.51 [11.81 33.24] | 19.07±7.22 [11.40 33.86] | .928 | 17/3 | 14/4 | .566 | 110.15±9.10 [97.00 130.00] | 114.00±12.08 [96.00 131.00] | .272 |
| SDSU | 16 | 7 | 1/6/0 | 13.98±1.94 [8.67 16.88] | 15.31±1.89 [12.13 17.15] | .143 | 10/6 | 7/0 | .059 | 106.94±10.99 [88.00 123.00] | 122.71±12.45 [112.00 141.00] | .006 |
| TRINITY | 23 | 19 | 7/6/6 | 17.48±3.66 [12.04 25.66] | 16.63±3.02 [12.00 23.08] | .424 | 23/0 | 19/0 | -- | 110.65±12.15 [89.00 133.00] | 110.63±14.10 [89.00 135.00] | .996 |
| UCLA_1 | 25 | 24 | 24/0/0 | 13.55±1.99 [9.50 17.79] | 13.48±2.72 [8.49 17.94] | .914 | 21/4 | 22/2 | .413 | 104.12±9.69 [84.00 126.00] | 104.96±12.19 [86.00 132.00] | .791 |
| UCLA_2 | 9 | 7 | 7/0/0 | 12.23±1.14 [9.79 13.63] | 12.68±1.99 [10.57 16.47] | .576 | 7/2 | 7/0 | .182 | 112.44±10.36 [99.00 128.00] | 94.71±11.63 [86.00 118.00] | .006 |
| UM_1 | 34 | 18 | 14/3/1 | 14.02±3.14 [9.50 19.20] | 13.60±2.44 [9.70 18.60] | .621 | 22/12 | 13/5 | .583 | 109.34±9.64 [89.00 127.50] | 107.81±12.32 [89.50 135.00] | .623 |
| UM_2 | 17 | 10 | 8/2/0 | 17.14±4.27 [13.60 28.80] | 15.28±1.49 [13.10 17.40] | .198 | 16/1 | 9/1 | .693 | 109.88±10.10 [89.50 129.00] | 113.25±14.73 [90.50 133.50] | .487 |
| Total | 203 | 133 | 93/33/7 | 16.66±6.33 [7.00 46.00] | 17.25±7.76 [8.09 8.09] | .447 | 167/36 | 118/15 | .107 | 109.92±10.43 [84.00 146.00] | 109.33±13.78 [84.00 141.00] | .655 |

Table S6 (continued).

| | ADOS, mean±SD [Range] | | | |
|---|---|---|---|---|
| Site | ADOS_TOTAL | ADOS_COMM | ADOS_SOCIAL | ADOS_STEREO |
| KKI | 12.20±3.19 [8.00 16.00] | 3.20±0.84 [2.00 4.00] | 9.00±2.74 [6.00 13.00] | 3.40±2.30 [0.00 6.00] |
| LEUVEN_1 | -- | -- | -- | -- |
| MAX_MUN | -- | -- | -- | -- |
| PITT | 12.50±3.71 [7.00 19.00] | 4.19±1.33 [2.00 7.00] | 8.31±2.73 [5.00 12.00] | 2.08±1.55 [0.00 6.00] |
| SDSU | -- | -- | -- | -- |
| TRINITY | -- | -- | -- | -- |
| UCLA_1 | 10.42±4.21 [2.00 17.00] | 3.25±1.65 [0.00 6.00] | 7.17±2.87 [2.00 11.00] | 1.33±1.69 [0.00 7.00] |
| UCLA_2 | 13.50±2.81 [9.00 16.00] | 3.67±1.51 [1.00 5.00] | 9.83±1.60 [8.00 12.00] | 3.00±1.79 [1.00 6.00] |
| UM_1 | -- | -- | -- | -- |
| UM_2 | -- | -- | -- | -- |
| Total[c] | 11.61±3.91 [2.00 19.00] | 3.59±1.50 [0.00 7.00] | 8.02±2.78 [2.00 13.00] | 1.96±1.83 [0.00 7.00] |

Abbreviations. TDC, typically developing control; ADOS, Autism Diagnostic Observation Schedule; ADOS_TOTAL, classic total ADOS score; ADOS_COMM, communication total sub-score of the classic ADOS; ADOS_SOCIAL, social total score of the classic ADOS; ADOS_STEREO, stereotyped behaviors and restricted interests total sub-score of the classic ADOS. KKI, Kennedy Krieger Institute; LEUVEN_1, University of Leuven: Sample 1; MAX_MUN, Ludwig Maximilians University Munich; PITT, University of Pittsburgh School of Medicine; SDSU, San Diego State University;

TRINITY, Trinity Centre for Health Sciences; UCLA_1, University of California Los Angeles: Sample 1; UCLA_2, University of California Los Angeles: Sample 2; UM_1, University of Michigan: Sample 1; UM_2, University of Michigan: Sample 2.

a, two sample Student's *t*-test with two tails; b, $\chi^2$ test; c, 51 of 126 ASD subjects have ADOS_TOTAL, ADOS_COMM, ADOS_SOCAIL data available and 48 for ADOS_STRERO; "--" indicates data unavailable.

**Table S7. Characteristics of human ABIDE-II cohorts.**

| Site | Number | | Autism/Asperger/PDD-NOS | Age, mean±SD [Range] | | | Gender (M/F) | | | FIQ, mean±SD [Range] | | |
|---|---|---|---|---|---|---|---|---|---|---|---|---|
| | HC | ASD | | HC | ASD | $p^a$ | HC | ASD | $p^b$ | HC | ASD | $p^a$ |
| GU_1 | 39 | 23 | 9/12/2 | 10.49±1.74<br>[8.06 13.80] | 11.29±1.05<br>[9.22 13.13] | .051 | 18/21 | 20/3 | .001 | 121.08±14.05<br>[95.00 149.00] | 117.61±13.98<br>[92.00 139.00] | .351 |
| NYU_1 | 22 | 17 | 5/0/12 | 9.68±3.78)<br>[5.90 23.81] | 10.43±6.94<br>[5.43 34.76] | .669 | 20/2 | 16/1 | .709 | 116.27±14.48<br>[91.00 144.00] | 111.88±14.79<br>[87.00 138.00] | .358 |
| TCD_1 | 18 | 12 | 3/9/0 | 16.28±2.79<br>[12.00 20.00] | 14.75±3.54<br>[10.00 19.50] | .198 | 18/0 | 12/0 | -- | 120.11±10.60<br>[99.00 142.00] | 113.92±14.01<br>[83.00 139.00] | .179 |
| UCLA_1 | 10 | 8 | 8/0/0 | 9.93±2.15<br>[7.76 14.09] | 12.85±1.80<br>[9.66 15.03] | .007 | 6/4 | 8/0 | .043 | 117.80±14.14<br>[94.00 141.00] | 106.25±9.95<br>[93.00 118.00] | .069 |
| Total | 89 | 60 | 25/21/14 | 11.40±3.59<br>[5.90 23.81] | 11.95±4.33<br>[5.43 5.43] | .402 | 62/27 | 56/4 | .000 | 119.33±13.47<br>[91.00 149.00] | 113.73±13.96<br>[83.00 139.00] | .016 |

Table S7 (continued).

| Site | ADOS_G, mean±SD [Range] | | | |
|---|---|---|---|---|
| | ADOS_TOTAL | ADOS_COMM | ADOS_SOCIAL | ADOS_STEREO |
| GU_1 | 10.53±4.79<br>[3.00 18.00] | 3.00±1.45<br>[1.00 7.00] | 7.53±3.81<br>[2.00 14.00] | 1.79±1.72<br>[0.00 5.00] |
| NYU_1 | 8.64±2.21<br>[5.00 12.00] | 2.14±1.10<br>[1.00 4.00] | 6.50±1.29<br>[4.00 8.00] | 1.36±1.01<br>[0.00 4.00] |
| TCD_1 | 8.25±1.82<br>[7.00 12.00] | 2.75±0.75<br>[2.00 4.00] | 5.50±1.62<br>[3.00 8.00] | 0.17±0.58<br>[0.00 2.00] |
| UCLA_1 | 13.00±0.71<br>[12.00 14.00] | 3.80±0.84<br>[3.00 5.00] | 9.20±0.45<br>[9.00 10.00] | 3.20±1.64<br>[1.00 5.00] |
| Total[c] | 9.70±3.56<br>[3.00 18.00] | 2.78±1.23<br>[1.00 7.00] | 6.92±2.75<br>[2.00 14.00] | 1.42±1.55<br>[0.00 5.00] |

Abbreviations. TDC, typically developing control; ADOS, Autism Diagnostic Observation Schedule; ADOS_TOTAL, classic total ADOS score; ADOS_COMM, communication total sub-score of the classic ADOS; ADOS_SOCIAL, social total score of the classic ADOS; ADOS_STEREO, stereotyped behaviors and restricted interests total sub-score of the classic ADOS. GU_1, Georgetown University; NYU_1, New York University Langone Medical Center: Sample 1; TCD_1, Trinity Centre for Health Sciences; UCLA_1, University of California Los Angeles.

a, two sample Student's *t*-test with two tails; b, $\chi^2$ test; c, 48 of 60 ASD subjects have ADOS data available; "--" indicates data unavailable.

**Table S8. Characteristics of human OCD cohorts.**

| Site | Number | | Age, mean±SD [Range] | | | Gender (M/F) | | | OCD, mean±SD | |
|---|---|---|---|---|---|---|---|---|---|---|
| | HC | OCD | HC | OCD | $p^a$ | HC | OCD | $p^b$ | Y-BOCS | HAM-A |
| ION | 58 | 78 | 30.79±8.41 [21.00 62.00] | 31.04±8.94 [14.00 63.00] | .871 | 38/20 | 45/33 | .355 | 30.12±6.77 | 17.92±10.27 |
| Ruijin | 21 | 14 | 30.81±5.84 [24.00 45.00] | 27.29±10.87 [16.00 46.00] | .221 | 13/8 | 10/4 | .561 | 24.29±5.50 | 15.50±9.67 |
| Total | 79 | 92 | 30.80±7.77 [21.00 62.00] | 30.47±9.30 [14.00 14.00] | .803 | 51/28 | 55/37 | .521 | 29.23±6.89 | 17.55±10.17 |

Abbreviations. OCD, Obsessive compulsive disorder; Y-BOCS, Yale-Brown Obsessive Compulsive Scale; HAM-A, Hamilton Anxiety Rating Scale.

a, two sample Student's *t*-test with two tails; b, $\chi^2$ test.

**Table S9. Characteristics of human ADHD cohorts.**

| Site | Number | | Age, mean±SD [Range] | | | Gender (M/F) | | | FIQ, mean±SD) [Range] | | |
|---|---|---|---|---|---|---|---|---|---|---|---|
| | HC | ADHD | HC | ADHD | p-value[a] | HC | ADHD | p-value[b] | HC | ADHD | p-value[a] |
| KKI | 47 | 16 | 10.44±1.30<br>[8.12 12.87] | 10.60±1.63<br>[8.10 12.99] | 0.684 | 27/20 | 8/8 | 0.605 | 109.70±11.48<br>[85.00 134.00] | 110.31±14.01<br>[88.00 134.00] | 0.863 |
| OHSU | 26 | 18 | 9.20±1.26<br>[7.33 11.92] | 9.09±1.15<br>[7.42 11.83] | 0.784 | 13/13 | 13/5 | 0.14 | 119.19±13.11<br>[98.00 144.00] | 107.00±13.88<br>[82.00 132.00] | 0.005 |
| Peking_1 | 57 | 20 | 11.12±1.61<br>[8.42 14.83] | 11.35±2.35<br>[9.00 17.33] | 0.627 | 16/41 | 16/4 | 0.000 | 118.23±13.85<br>[81.00 143.00] | 97.85±12.40<br>[81.00 128.00] | 0.000 |
| Peking_2 | 22 | 31 | 11.53±1.85<br>[9.08 14.33] | 12.66±1.74<br>[9.25 15.83] | 0.028 | 22/0 | 31/0 | -- | 121.45±13.68<br>[94.00 153.00] | 112.03±12.33<br>[86.00 135.00] | 0.012 |
| Peking_3 | 21 | 17 | 13.24±0.99<br>[11.25 14.92] | 13.37±1.30<br>[11.00 16.00] | 0.738 | 21/0 | 17/0 | -- | 113.14±13.05<br>[84.00 135.00] | 101.59±10.28<br>[83.00 120.00] | 0.005 |
| Total | 173 | 102 | 10.96±1.81<br>[7.33 14.92] | 11.57±2.23<br>[7.42 7.42] | 0.014 | 99/74 | 85/17 | 0.000 | 115.85±13.57<br>[81.00 153.00] | 106.35±13.53<br>[81.00 135.00] | 0.000 |

Abbreviations. TDC, typically developing control. KKI, Kennedy Krieger Institute; OHSU, Oregon Health & Science University; Peking, Peking University.

a, two sample Student's *t*-test with two tails; b, $\chi^2$ test; "--" indicates data unavailable.

**Table S10. Imaging protocols for resting-state fMRI used in the present study.**

ABIDE-I ASD cohort.

| Parameter | Site | | | | | | | | | |
|---|---|---|---|---|---|---|---|---|---|---|
| | KKI | LEUVEN_1 | MAX_MUN | PITT | SDSU | TRINITY | UCLA_1 | UCLA_2 | UM_1 | UM_2 |
| MRI Scanner | Philips Achieva | Philips | Siemens Magnetom Verio | Siemens Magnetom Allegra | GE MR750 | Philips Achieva | Siemens Magnetom TrioTim | Siemens Magnetom TrioTim | GE Signa | GE Signa |
| Magnetic field strength (T) | 3 | 3 | 3 | 3 | 3 | 3 | 3 | 3 | 3 | 3 |
| Field of view (mm) | 256 | 230 | 192 | 200 | 220 | 240 | 192 | 192 | 220 | 220 |
| Matrix | 84×81 | 64×64 | -- | 64×64 | 64×64 | 80×80 | 64×64 | 64×64 | | |
| Number of slices | 47 | 32 | 28 | 29 | | 38 | 34 | 34 | 40 | 40 |
| In-plane resolution (mm) | 3.05×3.15 | 3.59×3.59 | 3.0×3.0 | 3.1×3.1 | 3.4×3.4 | 3.0×3.0 | 3.0×3.0 | 3.0×3.0 | 3.4×3.4 | 3.4×3.4 |
| Slice thickness (mm) | 3 | 4 | 4 | 4 | 3.4 | 3.5 | 4 | 4 | 3 | 3 |
| Slice gap (mm) | 0 | 0 | -- | 0 | 0 | 0.35 | 0 | 0 | 0 | 0 |
| TR (ms) | 2500 | 1667 | 3000 | 1500 | 2000 | 2000 | 3000 | 3000 | 2000 | 2000 |
| TE (ms) | 30 | 33 | 30 | 25 | 30 | 28 | 28 | 28 | 30 | 30 |
| Total scan time (mm:ss) | 6:40 | 7:06 | 6:06 | 5:06 | 6:10 | 5:06 | 6:06 | 6:06 | 10:00 | 10:00 |
| Flip angle | 75 | 90 | 80 | 70 | 90 | 90 | 90 | 90 | 90 | 90 |
| Slice acquisition order | Ascending | Ascending | -- | Ascending | Ascending | Ascending | Ascending | Ascending | -- | -- |
| Eyes during scan | Opened | Opened | Closed/Opened | Closed | Opened | Closed | Opened | Opened | Opened | Opened |

KKI, Kennedy Krieger Institute; LEUVEN_1, University of Leuven: Sample 1; MAX_MUN, Ludwig Maximilians University Munich; PITT, University of Pittsburgh School of Medicine; SDSU, San Diego State University; TRINITY, Trinity Centre for Health Sciences; UCLA_1, University of California Los Angeles: Sample 1; UCLA_2, University of California Los Angeles: Sample 2; UM_1, University of Michigan: Sample 1; UM_2, University of Michigan: Sample 2.

Table S10 (continued).

ABIDE-II ASD cohort

| Parameter | Site | | | |
|---|---|---|---|---|
| | GU_1 | NYU_1 | TCD_1 | UCLA_1 |
| MRI Scanner | Siemens Trio | Siemens Allegra | Philips Achieva | Siemens Magnetom TrioTim |
| Magnetic field strength (T) | 3 | 3 | 3 | 3 |
| Field of view (mm) | 192 | 192 | 240 | 192 |
| Matrix | 64×64 | 64×64 | 80×80 | 64×64 |
| Number of slices | 43 | 33 | 37 | 34 |
| In-plane resolution (mm) | 3.0×3.0 | 3.0×3.0 | 3.0×3.0 | 3.0×3.0 |
| Slice thickness (mm) | 2.5 | 3 | 3.2 | 4 |
| Slice gap (mm) | 0.5 | 0 | 0.3 | 0 |
| TR (ms) | 2000 | 2000 | 2000 | 3000 |
| TE (ms) | 30 | 15 | 27 | 28 |
| Total scan time (mm:ss) | 5:14 | 6:00 | 7:06 | 6:06 |
| Flip angle | 90 | 82 | 90 | 90 |
| Slice acquisition order | Ascending | Ascending | Ascending | Ascending |
| Eyes during scan | Opened | Opened | Opened | Opened |

GU_1, Georgetown University; NYU_1, New York University Langone Medical Center: Sample 1; TCD_1, Trinity Centre for Health Sciences; UCLA_1, University of California Los Angeles.

ADHD cohort

| Parameter | Site | | | | |
|---|---|---|---|---|---|
| | KKI | OHSU | Peking_1 | Peking_2 | Peking_3 |
| MRI Scanner | Siemens Trio | Siemens Magnetom TrioTim | Siemens Magnetom TrioTim | Siemens Magnetom TrioTim | Siemens Magnetom TrioTim |
| Magnetic field strength (T) | 3 | 3 | 3 | 3 | 3 |
| Field of view (mm) | 256 | 240 | 200 | 200 | 200 |
| Matrix | 84×81 | -- | -- | -- | -- |
| Number of slices | 47 | 36 | 33 | 33 | 33 |
| In-plane resolution (mm) | 3.05×3.15 | 3.8×3.8 | 3.1×3.1 | 3.1×3.1 | 3.1×3.1 |
| Slice thickness (mm) | 3 | 3.8 | 3.5 | 3.5 | 3.5 |
| Slice gap (mm) | 0 | -- | -- | -- | -- |
| TR (ms) | 2500 | 2000 | 2000 | 2000 | 2000 |
| TE (ms) | 30 | 30 | 30 | 30 | 30 |
| Total scan time (mm:ss) | 6:40 | 3:32 | 8:06 | 8:06 | 8:06 |
| Flip angle | 75 | 90 | 90 | 90 | 90 |
| Slice acquisition order | Ascending | Ascending | Ascending | Ascending | Ascending |
| Eyes during scan | Opened | Opened | Opened/Closed | Opened/Closed | Opened/Closed |

KKI, Kennedy Krieger Institute; OHSU, Oregon Health & Science University; Peking, Peking University
